# *ACTN2* mutant causes proteopathy in human iPSC-derived cardiomyocytes

**DOI:** 10.1101/2021.10.28.466251

**Authors:** Antonia T. L. Zech, Maksymilian Prondzynski, Sonia R. Singh, Ellen Orthey, Erda Alizoti, Josefine Busch, Alexandra Madsen, Charlotta S. Behrens, Giulia Mearini, Marc D. Lemoine, Elisabeth Krämer, Diogo Mosqueira, Sanamjeet Virdi, Daniela Indenbirken, Maren Depke, Manuela Gesell Salazar, Uwe Völker, Ingke Braren, William T. Pu, Thomas Eschenhagen, Elke Hammer, Saskia Schlossarek, Lucie Carrier

## Abstract

Genetic variants in α-actinin-2 (ACTN2) are associated with several forms of (cardio)myopathy. We previously reported a heterozygous missense (c.740C>T) *ACTN2* gene variant, associated with hypertrophic cardiomyopathy, and characterized by an electro-mechanical phenotype in human induced pluripotent stem cell-derived cardiomyocytes (hiPSC-CMs). Here, we created with CRISPR/Cas9 genetic tools two heterozygous functional knock-out hiPSC lines with a second wild-type (ACTN2wt) and missense ACTN2 (ACTN2mut) allele, respectively. We evaluated their impact on cardiomyocyte structure and function, using a combination of different technologies, including immunofluorescence and live cell imaging, RNA-seq, and mass spectrometry. This study showed that ACTN2mut present a higher percentage of multinucleation, protein aggregation, hypertrophy, myofibrillar disarray and activation of both the ubiquitin-proteasome system and the autophagy-lysosomal pathway as compared to ACTN2wt in 2D-cultured hiPSC-CMs. Furthermore, the expression of ACTN2mut was associated with a marked reduction of sarcomere-associated protein levels in 2D-cultured hiPSC-CMs and force impairment in engineered heart tissues. In conclusion, our study highlights the activation of proteolytic systems in ACTN2mut hiPSC-CMs likely to cope with ACTN2 aggregation and therefore directs towards proteopathy as an additional cellular pathology caused by this *ACTN2* variant, which may contribute to human *ACTN2*-associated cardiomyopathies.

## Introduction

α-Actinin-2 (ACTN2) is a component of the sarcomere in skeletal and cardiac myocytes, which forms anti-parallel homodimers that can anchor and crosslink actin thin filaments to the Z-disk (for reviews, see [1,2]). Additionally, ACTN2 is implicated in assembling large protein complexes for structural integrity, mechanotransduction and cell signaling (for reviews, see [3,4]).

Genetic variants in *ACTN2* are associated in the heterozygous state with common inherited cardiac diseases, which are hypertrophic (HCM), dilated (DCM), and restrictive (RCM) cardiomyopathy [5-8]. Although considered as a rare disease gene [9], recent reports showed the association of homozygous *ACTN2* variants with core myopathy [10], or progressive, severe RCM [6]. In addition, it was shown that heterozygous *ACTN2* variants have critical effects on structure and function of the cardiac muscle [11]. We previously demonstrated that a heterozygous HCM missense *ACTN2* variant (c.740C>T) induced an electro-mechanical phenotype in human induced pluripotent stem cell-derived cardiomyocytes (hiPSC-CMs; [8]). The molecular mechanisms by which missense *ACTN2* variants lead to different forms of (cardio)myopathy are not fully understood. It is assumed that mutant transcripts are translated into proteins, which are expected to have a dominant-negative effect on sarcomere structure and/or function. However, mutant proteins can also be misfolded and targeted towards the ubiquitin-proteasome system (UPS) for degradation or can form aggregates, causing cellular proteopathy if not targeted for degradation towards the autophagy-lysosomal pathway (ALP; [12-14]).

In this study, we created with CRISPR/Cas9 genetic tools two cell lines expressing either a wild-type ACTN2 (ACTN2wt; c.740C) or a mutant ACTN2 (ACTN2mut; c.740T) and evaluated the impact of the mutant ACTN2 on cellular structure and function in hiPSC-CMs, using a combination of different technologies, including immunofluorescence and live cell imaging, RNA-seq and mass spectrometry (MS) analyses. Our data showed that the ACTN2mut hiPSC-CMs present hypertrophy, myofibrillar disarray, ACTN2 aggregation, a higher percentage of multinucleation, and activation of both UPS and ALP. It was associated with a marked reduction of sarcomere-associated protein levels and force impairment in engineered heart tissues (EHTs). Our data indicate proteopathy as an additional cellular feature caused by the missense *ACTN2* variant, which may contribute to human *ACTN2*-associated cardiomyopathy.

## 2. Materials and Methods

### 2.1. Generation and culture of hiPSC-CMs in 2D and EHT formats

Cultivation, CRISPR/Cas9 gene editing, and differentiation of hiPSC lines into CMs were as previously described [8]. A detailed methodology is provided in the Supplement.

### 2.2. Morphological analysis of 2D-cultured hiPSC-CMs

Quantification of myofibrillar disarray and cell area was evaluated with Fiji (ImageJ) as described previously [8,15]. Detailed information for ACTN2 protein aggregate analysis and hiPSC-CMs volume measurement is provided in the Supplement.

### 2.3. Production and purification of adeno-associated virus vector particles

The production and purification of adeno-associated virus serotype 6 (AAV6) vector particles carrying the mTagRFP-mWasabi-hLC3 tandem construct and the WT- or MUT-*ACTN2*-HaloTag® was adapted from a recent publication [16]. A detailed protocol is provided in the Supplement.

### 2.4. Proteome analysis

Sample preparation (n = pool of 2-3 replicates per batch from 3 independent differentiation batches), protein digestion, and liquid chromatography–tandem MS were performed as described previously [17,18]. Detailed information is provided in the Supplement.

### 2.5. High-content imaging of autophagy-lysosomal pathway in hiPSC-CMs

A high-content screen for ALP activity was performed in 2D-cultured hiPSC-CMs transduced with an AAV6 encoding the mTagRFP-mWasabi-hLC3 tandem construct under the control of human *TNNT2* promoter. After 30 days of culture, hiPSC-CMs were fixed, stained with a cardiac troponin T antibody (TNNT2, 1:500; ab45932) and with Hoechst 33342 for nuclei staining (1 μg/mL; Thermo Fisher Scientific, Waltham, MA, USA), and imaged with the Operetta high-content imaging system (PerkinElmer, Nottingham, UK). Subsequent image analysis was performed with the Harmony high-content imaging analysis software (PerkinElmer, Nottingham, UK) by identifying TNNT2+ cells and quantifying green and red puncta (number and intensity). A detailed method is described in the Supplement.

### 2.6. Statistics

Group data are presented as mean ± SEM. GraphPad Prism 9 (GraphPad Software, San Diego, CA, USA) was used for data analysis. Curves were fitted to data points from individual experiments. When two groups were analyzed, data were compared with the unpaired Student’s t-test, nested t-tests when appropriate or two-way ANOVA followed by Tukey’s post-test, as described in the figure legends. Chi-square analysis was performed for multinucleation and integration/aggregation analysis, whereby each data set was compared to ACTN2wt using “compare observed distribution with expected” and the Wilson/Brown method. A mixed-effects analysis plus Sidak’s post-test was performed when pooled batches of two cell lines were analyzed over a timeline. A *P*-value < 0.05 was considered as statistically significant.

## 3. Results

## 3.1. ACTN2mut 2D-cultured hiPSC-CMs display hypertrophy, myofibrillar disarray, protein aggregation and multinucleation

Both ACTN2wt and ACTN2mut lines were derived from the previously described heterozygous ACTN2 HCM hiPSC line (ACTN2het; c.740C>T; [8]) using CRISPR/Cas9 gene editing and homology directed repair. Initially, both hiPSC lines were intended to produce an isogenic control without the mutation. However, sequencing analysis revealed only one functional wild-type allele in ACTN2wt [19] and only one functional mutant allele in ACTN2mut (Figure S1A). The other allele of both cell lines contained on-target defects of CRISPR/Cas9 (splice-site mutation in ACTN2wt [19] and large rearrangement in ACTN2mut (Figure S1A,B)), leading to nonsense mRNAs, which are not visible by RNA-seq (Figure S2D). Thus, we created two heterozygous functional knock-out lines with a second wild-type and missense ACTN2 allele, respectively. Both ACTN2wt and ACTN2mut hiPSC lines presented a normal karyotype (Figure S1C) and were therefore differentiated to hiPSC-CMs according to our protocol (Figure S1D). HiPSC-CMs were produced with high purity (on average >90% cardiac troponin T (TNNT2)-positive cells) and quantity (Figure S2A).

Both hiPSC-CM lines were evaluated for ACTN2 abundance and localization, myofibrillar disarray, cellular hypertrophy and multinucleation. Immunofluorescence analysis revealed a cross-striated pattern of TNNT2 in both hiPSC-CM lines, indicating proper formation of sarcomeres (Figure 1A). However, ACTN2 was less organized and formed aggregates in ACTN2mut when compared to ACTN2wt hiPSC-CMs. Quantification revealed a higher index of myofibrillar disarray and more ACTN2 aggregates in ACTN2mut hiPSC-CMs (Figure 1B,C). Further, cell area and volume were higher in ACTN2mut hiPSC-CMs (Figure 1D,E). Finally, the percentage of multinucleated (>1 nucleus) hiPSC-CMs was higher in ACTN2mut than in ACTN2wt (Figure 1F). Mononucleated-to-multinucleated ratio was 80:20 in ACTN2wt hiPSC-CMs, supporting previous estimation obtained in human hearts [20], whereas it was 53:47 in ACTN2mut, as reported in other HCM hiPSC-CMs [21]. *ACTN2* mRNA level did not differ between the two hiPSC-CM lines (Figure S2C,D), whereas ACTN2 protein level was markedly lower in ACTN2mut hiPSC-CMs (Figure S2E,F).

**Figure 1.**
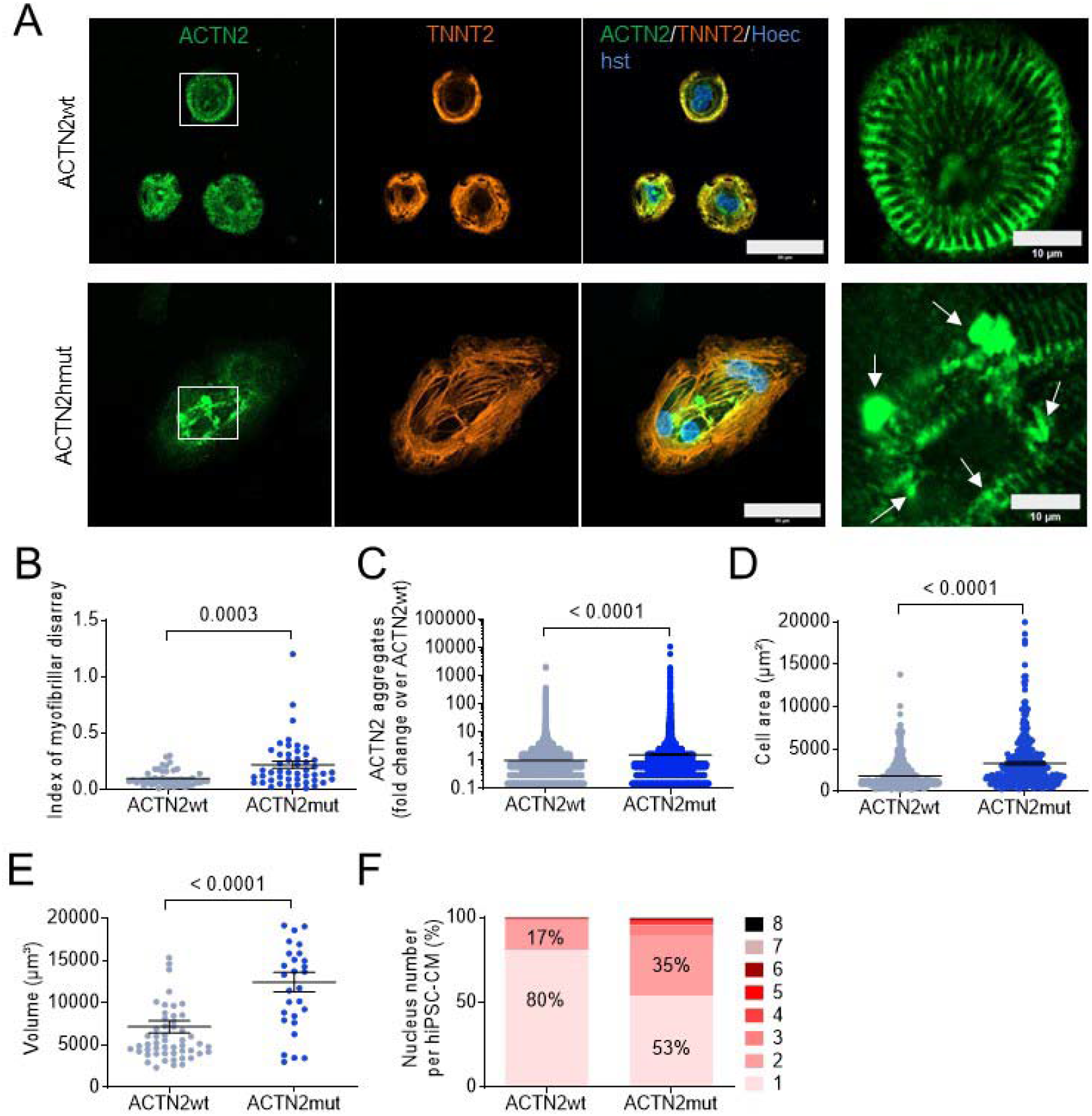
Disease modeling in 30-day-old, 2D-cultured hiPSC-CMs. **(A)** Representative immunofluorescence images of hiPSC-CMs (scale bar = 50 μm), including a higher magnification on the right (scale bar = 10 μm). After 30 days, hiPSC-CMs were fixed and stained with antibodies against ACTN2 and TNNT2, and with Hoechst for nuclei. **(B)** Blinded analysis of myofibrillar disarray using high-resolution pictures (ACTN2wt and ACTN2mut: N/d = 16/3). **(C)** Quantification of ACTN2 aggregates analyzed with Fiji software (ACTN2wt: N/d = 14/3, ACTN2mut: N/d = 15/3). **(D)** Quantification of cell area analyzed with Fiji software (ACTN2wt: N/n/d = 548/3/3, ACTN2mut: N/n/d 319/3/3). **(E)** Quantification of cell volume analyzed with Imaris software (ACTN2wt: N/n/d = 54/3/3, ACTN2mut: N/n/d = 29/3/3). **(F)** Quantification of nucleus number per hiPSC-CM analyzed with Fiji software and expressed as percentage (ACTN2wt: N/n/d = 548/3/3, ACTN2mut: N/n/d = 319/3/3; Chi-square = 280.1, *P* < 0.0001). Data are expressed as mean ± SEM (panels B-E) or percentages (panel F), with *P*-values obtained with the unpaired Student’s t-test (panels B-E) or two-tailed Chi-square test (panel F). Abbreviations: TNNT2, cardiac troponin T; N/n/d, number of cells/wells/differentiations. Arrows point to aggregates.

### 3.2. Exogenous mutant ACTN2 causes aggregate formation leading to sarcomere disarray

We then tested whether exogenous mutant ACTN2 could induce protein aggregation in living, 2D-cultured ACTN2wt hiPSC-CMs. Therefore, ACTN2wt hiPSC-CMs were transduced with AAV6 carrying the MUT-*ACTN2*-HaloTag® (c.740T) and were compared to ACTN2mut hiPSC-CMs transduced with AAV6 carrying the WT-*ACTN2*-HaloTag® (c.740C). After 7 days of culture in 96-well plates, live-cell imaging experiments were performed by staining ACTN2-HaloTag protein using TMR-ligand in combination with Hoechst (Figure 2A; and examples in Videos 1, 2). Exogenous MUT-ACTN2 in ACTN2wt and WT-ACTN2 in ACTN2mut reversed the phenotypes, inducing aggregation in about 83% and 17% of hiPSC-CMs, respectively (Figure 2A,B). This indicates that MUT-ACTN2 causes aggregation in ACTN2wt and WT-ACTN2 reverses aggregation in ACTN2mut. Quantification of acquired live cell images for ACTN2 aggregates revealed more and larger aggregates in MUT-*ACTN2*-transduced ACTN2wt than in WT-*ACTN2*-transduced ACTN2mut (Figure 2C). Western blot analysis revealed only the endogenous ACTN2 in MUT-*ACTN2*-transduced ACTN2wt (Figure 2D,E), suggesting a very low level of correctly folded MUT-ACTN2, which should be visible as a larger molecular weight protein. Conversely, both exogenous and endogenous ACTN2 were detected in WT-*ACTN2*-transduced ACTN2mut hiPSC-CMs, with about 46% of replacement of endogenous by exogenous ACTN2 (Figure 2F,G).

**Figure 2.**
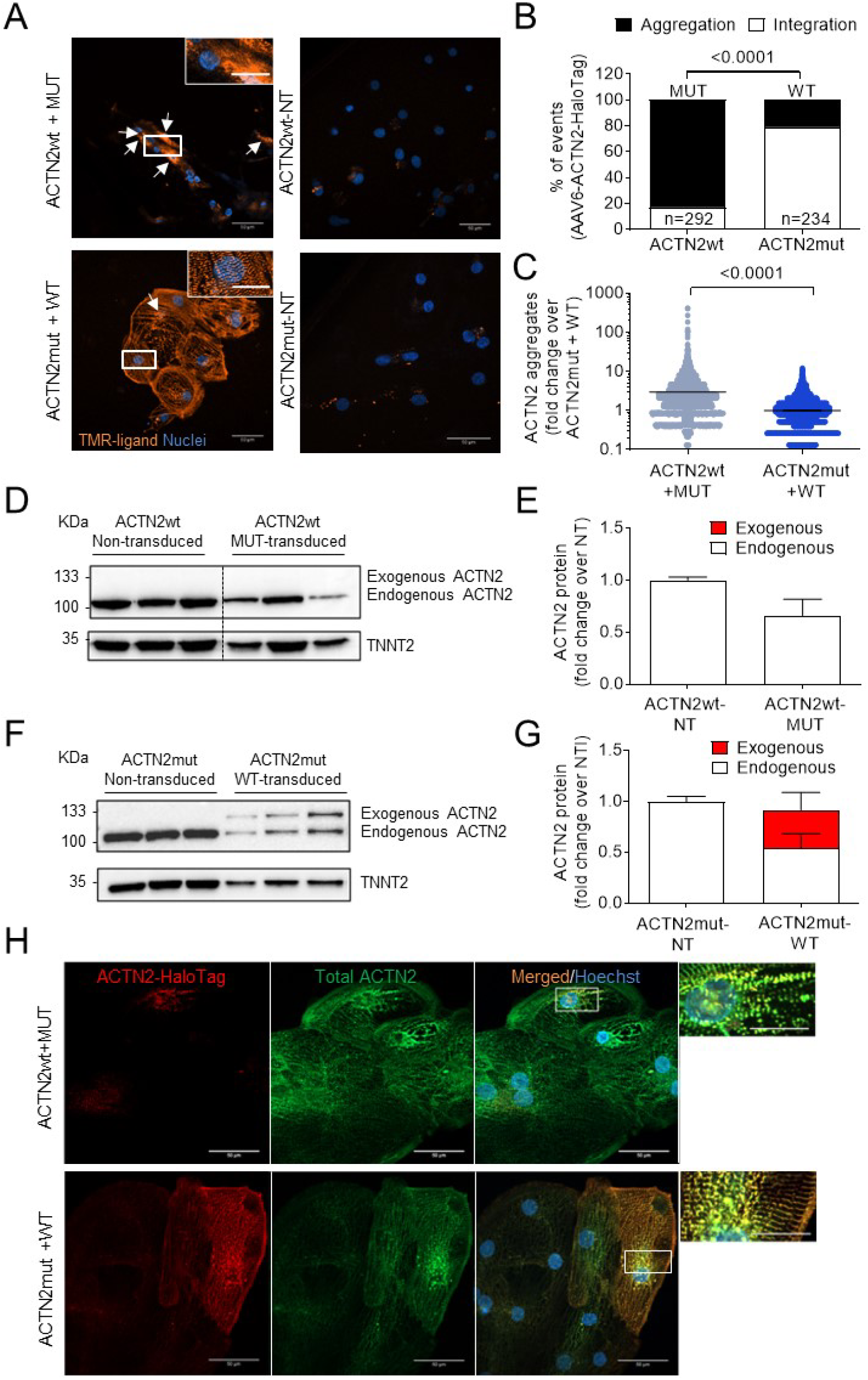
Live cell imaging and immunofluorescence in 2D-cultured hiPSC-CMs after 7 or 30 days in culture. **(A)** Representative images of ACTN2wt and ACTN2mut hiPSC-CMs transduced with an AAV6 carrying either the *ACTN2*-MUT-HaloTag® or *ACTN2*-WT-HaloTag® construct and seeded in 96-well plates (2,500-5,000 cells/well). After 7 days, live cell imaging was performed by adding TMR-ligand to stain ACTN2-HaloTag® protein and Hoechst for nuclei staining (Scale bars = 50 μm; zoom = 20 μm). **(B)** Blinded quantification of sarcomere integration or aggregation of exogenous MUT-ACTN2 in ACTN2wt hiPSC-CMs (N/n/d = 292/3/1) or WT-ACTN2 in ACTN2mut hiPSC-CMs (N/n/d = 234/3/1). **(C)** Quantification of ACTN2 aggregates in hiPSC-CM lines. Analysis was performed with Fiji software (ACTN2wt+MUT-ACTN2: N/d = 26/1, ACTN2mut+WT-ACTN2: N/d = 9/1). **(D, E)** Western blot stained for ACTN2 and TNNT2 and quantification of non-transduced ACTN2wt or transduced with *ACTN2*-MUT-HaloTag® (n/d=3/1) after 30 days. **(F, G)** Western blot stained for ACTN2 and TNNT2 and quantification of non-transduced (NT) ACTN2mut or ACTN2mut transduced with *ACTN2*-WT-HaloTag® (n/d=3/1) after 30 days. **(H)** Representative images of fixed ACTN2wt and ACTN2mut hiPSC-CMs transduced with an AAV6 carrying either the *ACTN2*-MUT-HaloTag® or the *ACTN2*-WT-HaloTag® after 30 days of culture and stained with antibodies directed against the HaloTag® and ACTN2, and Hoechst for nuclei staining (scale bar = 50 μm). Images were taken with a Zeiss LSM 800 microscope. Data are expressed as percentages (panel B) or as mean ± SEM (panels C, E, G), with *P*-values obtained with one-tailed Chi-square test (panel B) or unpaired Student’s t-test (panel C). Abbreviations: AAV6, adeno-associated virus serotype 6; MUT, mutant; NT, non-transduced; TNNT2, cardiac troponin T; WT, wild-type; N/n/d, number of cells/wells/differentiations. Arrows point to aggregates.

Immunofluorescence analysis of hiPSC-CMs transduced with WT- or MUT-*ACTN2* using antibodies directed against the HaloTag® and total ACTN2 revealed co-localization of WT-ACTN2-HaloTag® and total ACTN2 staining in ACTN2mut, confirming Z-disk integration of exogenous WT-ACTN2, whereas exogenous MUT-ACTN2 was barely detectable in ACTN2wt and exhibited co-localization with total ACTN2 in some parts (Figure 2H).

### 3.3. ACTN2mut hiPSC-CMs exhibit alterations of several canonical pathways

To understand the molecular changes caused by the *ACTN2* mutation, MS was performed in 2D-cultured hiPSC-CMs. Three replicates of each hiPSC-CM line were pooled, and 3 batches of differentiation were analyzed. Volcano plots depict 481 (250 higher, 231 lower) dysregulated proteins in ACTN2mut vs. ACTN2wt (Figure 3A). Ingenuity Pathway Analysis (IPA) revealed dysregulation of several canonical pathways, diseases, and biological functions in ACTN2mut hiPSC-CMs (Figure 3B; Dataset S1). Specifically, mitochondrial function, sirtuin signaling, protein ubiquitination, hereditary myopathy, sliding of myofilaments and stabilization of mRNA were highly dysregulated in ACTN2mut. A deeper analysis of proteomic data revealed that the protein levels of ACTN2, several other sarcomere-associated proteins and desmosomal proteins were markedly lower in ACTN2mut hiPSC-CMs (Table S1). Some of these proteins (FLNC, MYOZ2, NEBL, SYNPO2, SYNPO2L, TTN) are known to interact directly with ACTN2 [22-24]. On the other hand, FHL1 and FHL2, located at the Z-disk of the sarcomere [25,26] were more abundant in ACTN2mut. Several proteins associated with the UPS and/or ALP (e.g. BAG3, CTSC, GBA, HSPA1A, PSMA3, PSMA6, PSMB5, PSME2, TRIM54, UBA1, UBE2O, UBQLN2) were more abundant in ACTN2mut hiPSC-CMs (Table S2).

**Figure 3.**
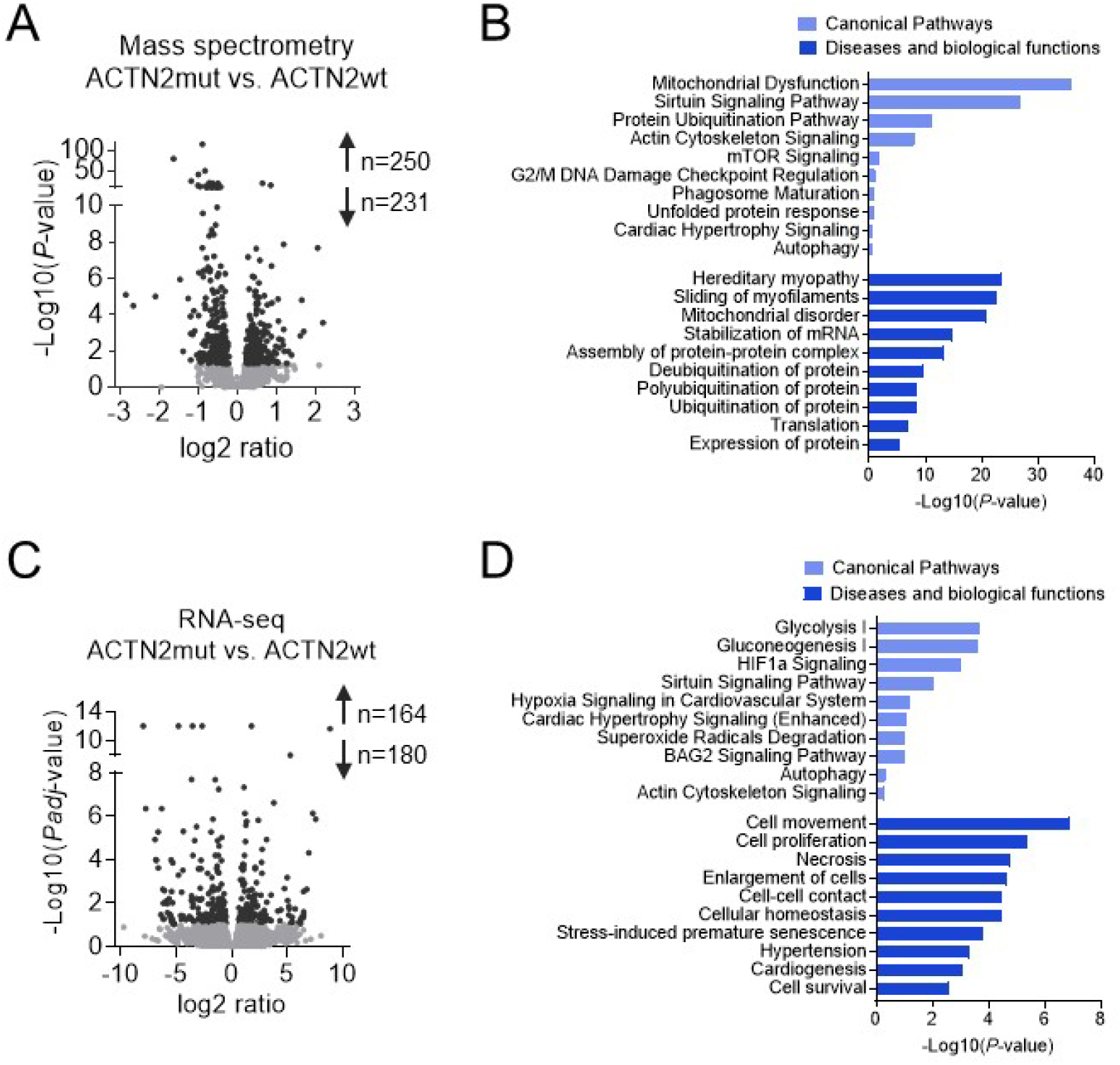
MS and RNA-seq analysis of 30-day-old, 2D-cultured hiPSC-CMs. **(A)** Alterations in protein levels between ACTN2mut vs. ACTN2wt (250 up, 231 down) hiPSC-CMs based on MS are displayed in a volcano plot that shows the -Log10 of *P*-value vs. magnitude of change (log2 ratio) whereby light grey dots indicate *P* > 0.05 and dark grey dots *P* < 0.05. **(B)** Selected hits of significantly dysregulated canonical pathways and biological functions in 2D-cultured ACTN2mut vs. ACTN2wt hiPSC-CMs based on MS analysis using Ingenuity Pathway Analysis (IPA). Unsupervised IPA was performed for significantly altered proteins (Fisher’s exact test; *P* < 0.05). **(C)** Alterations in mRNA levels in ACTN2mut vs. ACTN2wt (164 up, 180 down) hiPSC-CMs based on RNA-seq analysis are displayed in a volcano plot that shows the -Log10 of *Padj*-value vs. magnitude of change (log2 ratio) whereby light grey dots indicate FDR > 0.1 and dark grey dots FDR < 0.1. **(D)** Selected hits of significantly dysregulated canonical pathways and biological functions in 2D-cultured ACTN2mut vs. ACTN2wt hiPSC-CMs based on RNA-seq analysis using IPA. Unsupervised IPA was performed for significantly altered genes (Fisher’s exact test; FDR < 0.1).

We then performed RNA-seq on 3 pooled replicates of each hiPSC-CM line from 3 cardiac differentiation batches. Volcano plot showed 344 (164 higher, 180 lower) dysregulated mRNAs in ACTN2mut vs. ACTN2wt (Figure 3C). IPA analysis revealed several different dysregulated canonical pathways, diseases, and biological functions in ACTN2mut when compared to ACTN2wt (Figure 3D; Dataset S2). Some of the highlighted IPA pathways were metabolism, hypoxia, oxidative and cellular stress, cardiac hypertrophy, and cellular remodeling, whereas signaling of actin cytoskeleton was less pronounced. Specifically, the mRNA levels of sarcomere-associated proteins did not differ between the two groups, except for *FHL2* and *MYH6*, which were lower and higher in ACTN2mut than in ACTN2wt, respectively (Table S1). These data were confirmed by mRNA count analysis using the nanoString nCounter® Elements technology (Figure S2D,E). RNA-seq also revealed dysregulation of several genes, encoding proteins involved in the UPS and ALP in ACTN2mut hiPSC-CMs (Table S2).

Taken together, Omics analysis supported experimental findings for structural sarcomere abnormalities in ACTN2mut hiPSC-CMs and suggested alterations in pathways such as cellular stress response, cell survival/apoptosis or protein homeostasis, which directly point towards proteopathy as an important disease feature.

### 3.4. ACTN2mut hiPSC-CMs exhibit higher activities of the ubiquitin-proteasome system and the autophagy-lysosomal pathway

Higher abundance of several UPS- and/or ALP-associated proteins and presence of ACTN2 aggregates in ACTN2mut hiPSC-CMs suggested an altered proteostasis. Therefore, the activity of both systems was evaluated in 2D-cultured hiPSC-CMs. To evaluate the UPS, cells were treated either with vehicle (0.05% DMSO) or the UPS inhibitor epoxomicin (250 nM; Figure 4A-D). Under basal conditions (DMSO), the levels of (poly)ubiquitinated proteins and of their shuttle protein for autophagy-mediated degradation SQSTM1 did not differ between cell lines, whereas ACTN2 level was lower in ACTN2mut than ACTN2wt, reproducing our findings (Figure S2E,F). Epoxomicin treatment induced a marked accumulation of (poly)ubiquitinated proteins and SQSTM1 in both hiPSC-CMs (Figure 4A-C), validating the efficacy of the treatment. In contrast, epoxomicin did not increase the level of ACTN2 in any cell line, indicating that ACTN2 was not degraded by the UPS in this experimental condition (Figure 4A,D). On the other hand, the chymotrypsin-like activity of the proteasome was markedly higher in ACTN2mut hiPSC-CMs (Figure 4E), suggesting UPS activation.

**Figure 4.**
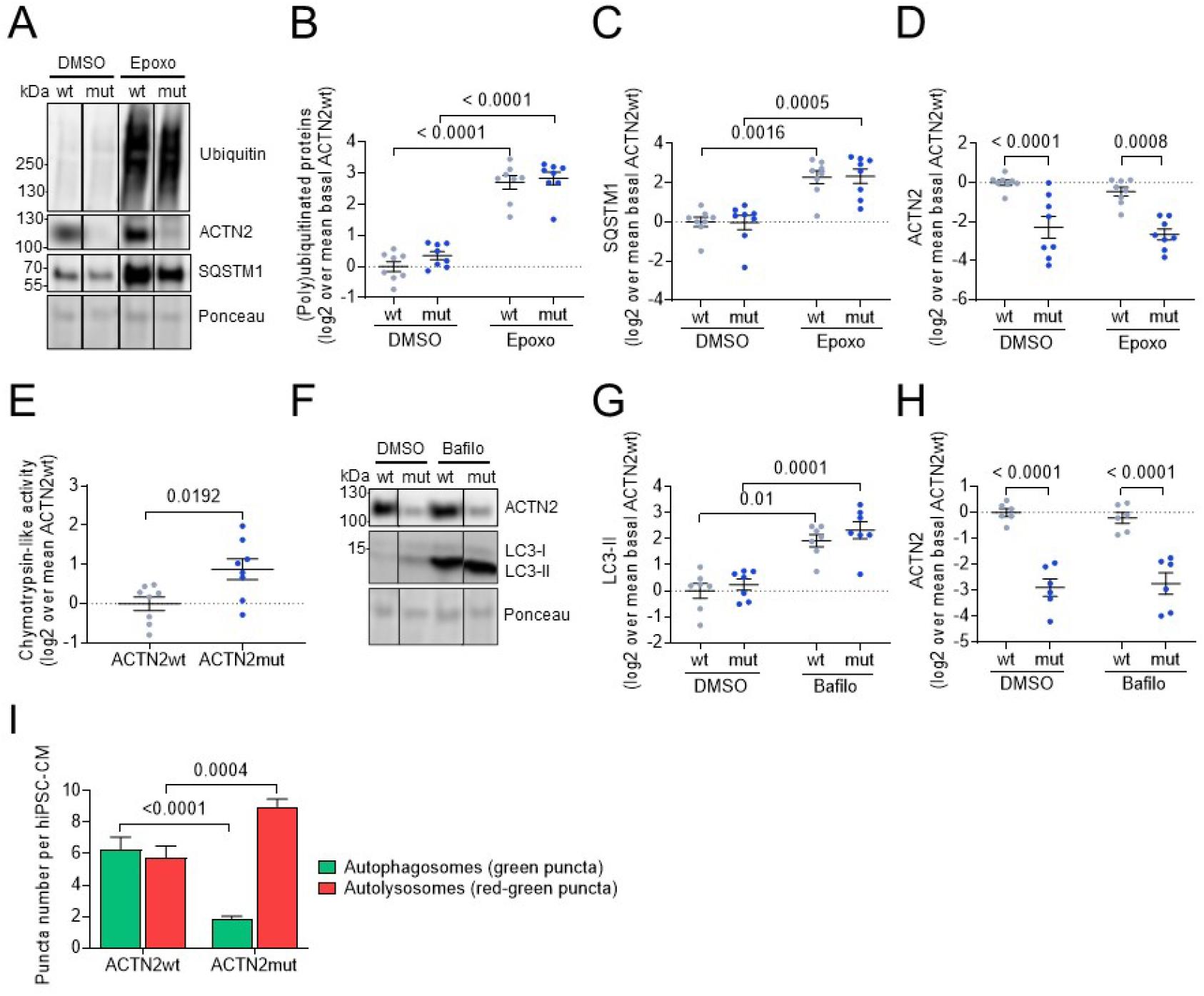
Evaluation of the impact of the *ACTN2* gene variant on the proteolytic systems in 30-day-old, 2D-cultured hiPSC-CMs. Representative Western blots and Ponceau. **(A)**, and quantification, normalized to Ponceau, of the levels of (B) (poly)ubiquitinated proteins, **(C)** SQSTM1, **(D)** ACTN2 in hiPSC-CMs treated with DMSO (0.05%) or epoxomicin (Epoxo; 250 nM) for 16.5 h at 37 °C (ACTN2wt: n/d = 7-8/3, ACTN2mut: n/d = 7-8/3). **(E)** Chymotrypsin-like activity of the proteasome in hiPSC-CMs (ACTN2wt: n/d = 8/3, ACTN2mut: n/d = 8/3). **(F)** Representative Western blots, Ponceau, and quantification of protein levels of **(G)** LC3-II and **(H)** ACTN2 of hiPSC-CMs treated with DMSO (0.05%) or bafilomycin A1 (Bafilo; 50 nM) for 3 h at 37 °C (ACTN2wt: n/d = 6-7/3, ACTN2mut: n/d = 6-7/3). **(I)** The ALP activity was indirectly measured by determining the amount of autophagosomes (green puncta) and autolysosomes (red minus green puncta) whereby the number of puncta is related to the CM number per well (ACTN2wt and ACTN2mut: n = 9). Data are expressed as mean ± SEM, with *P*-values obtained with two-way ANOVA and Tukey’s post-test (panels B-D, G-I) or with unpaired Student’s t-test (panels E). Abbreviations: n/d, number of wells/differentiations.

To evaluate the ALP, the autophagic flux was measured in hiPSC-CMs after treatment with either DMSO (0.05%) or the late-stage ALP inhibitor bafilomycin A1 (50 nM; Figure 4F-H). The level of microtubule-associated protein 1 light chain 3b-II (LC3-II) did not differ between the genotypes in basal conditions. Treatment with bafilomycin A1 markedly increased LC3-II levels in both groups (Figure 4F,G). The difference in LC3-II level between bafilomycin-treated and DMSO-treated samples, which represents the autophagic flux, was higher in ACTN2mut than in ACTN2wt (difference in log2, ACTN2wt: 1.93, ACTN2mut: 2.09). On the other hand, bafilomycin A1 did not increase ACTN2 levels (Figure 4H), implying that ACTN2 is not degraded by the ALP. To support the autophagic flux data, we performed a high-content imaging in the hiPSC-CM lines transduced with an AAV6 encoding mWasabi-mTagRFP-hLC3 under the control of the *TNNT2* promoter. After 30 days of culture, hiPSC-CMs were fixed and immunostained for TNNT2 and Hoechst to ensure imaging of (solely) cardiomyocytes (Figure S3A; >80% TNNT2+, data not shown). The number of green and red puncta per well was quantified using an unbiased and statistically powerful method and normalized to the number of hiPSC-CMs per well. The number of green puncta per hiPSC-CM was markedly lower in ACTN2mut (Figure S3B), whereas the number of red puncta per hiPSC-CM did not differ between the groups (Figure S3C). The utilization of the LC3-tandem-construct allows to determine autophagosomes (AP) and autolysosomes (AL), since the green fluorescence (mWasabi) is susceptible to low pH and hence quenched within ALs. Therefore, green puncta correspond to APs, red puncta to APs plus ALs, and the difference between red and green puncta (=red minus green puncta) to ALs. ACTN2wt hiPSC-CMs exhibited a similar number of APs and ALs per hiPSC-CM (Figure 4I), suggesting a steady-state autophagic flux. In contrast, the AP number per hiPSC-CM was markedly lower and the AL number per hiPSC-CM was higher in ACTN2mut (Figure 4I). The combination of low AP number and high AL number supports the view of an activation of autophagy, particularly at the step of fusion of APs with lysosomes to form autolysosomes in ACTN2mut hiPSC-CMs.

Taken together, these data showed higher activities of both protein degradation systems in 2D-cultured ACTN2mut hiPSC-CMs, most likely to eliminate protein aggregates causing proteopathy.

### 3.5. ACTN2mut hiPSC-CMs exhibit force impairment in engineered heart tissues

The low abundance of several sarcomere-associated proteins in ACTN2mut hiPSC-CMs (Table S1) suggested an impairment of contractile function. We therefore assessed force amplitude and kinetics of the ACTN2wt and ACTN2mut in 3D EHTs after 30 days (Figure 5A,B). Unpaced ACTN2mut EHTs developed significantly lower force starting from day 9 onwards than ACTN2wt EHTs (Figure 5C). Beating frequency was significantly higher in ACTN2mut than in ACTN2wt EHTs (50 vs. 28 beats per minute from day 21 on, respectively (Figure 5D). To compare functional parameters independent of variable baseline frequencies, EHTs were subjected to electrical pacing at 1 Hz (Videos 3, 4). Contraction traces of EHTs showed markedly lower force in ACTN2mut than ACTN2wt EHTs (Figure 5E,F). Normalized averaged force exhibited 19% shorter time to peak (TTP_-80%_; Figure 5G,H) and 25% shorter relaxation time (RT_80%_; Figure 5G,I) in ACTN2mut EHTs.

**Figure 5.**
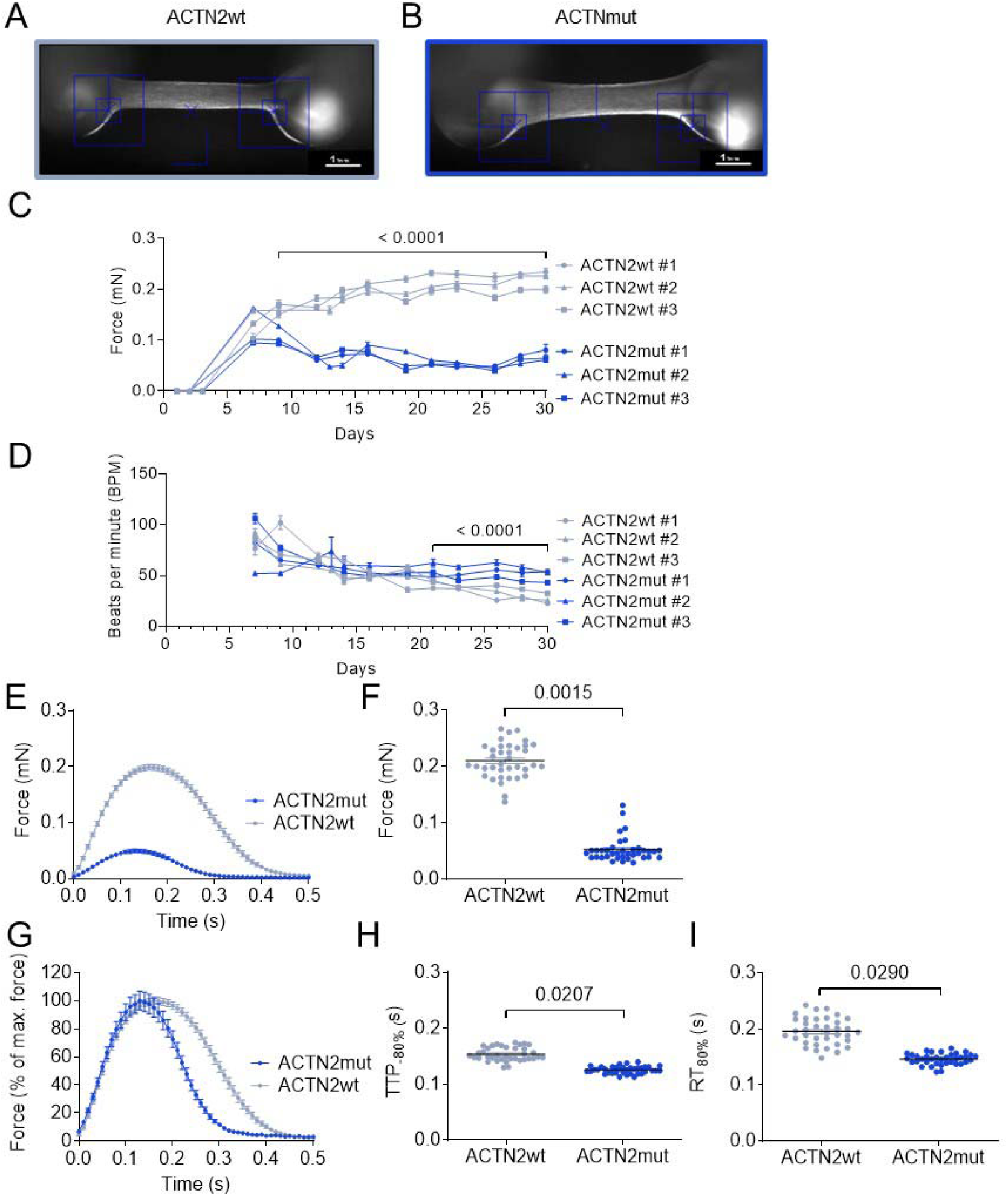
Force measurements in 3D-cultured hiPSC-engineered heart tissues. Representative images of **(A)** ACTN2wt and **(B)** ACTN2mut EHTs cultured for 30 days (scale bar = 1 mm). **(C)** Force and **(D)** beats per minute were measured under unpaced conditions in EHT culture medium or in 1.8 mM Ca^2+^ Tyrode’s solution at 37 °C (ACTN2wt: N/d = 27-44/3, ACTN2mut: N/d = 32-44/3). **(E)** Average force, **(F)** force, **(G)** normalized force, **(H)** time to peak -80% (TTP_-80%_), and **(I)** relaxation time to baseline 80% (RT_80%_) were measured under paced conditions at 1 Hz in 1.8 mM Ca^2+^ Tyrode’s solution at 37 °C (ACTN2wt: N/d = 37/3, ACTN2mut: N/d = 37/3); data are expressed as mean ± SEM, with *P*-values obtained with a mixed-effects analysis plus Sidak’s multiple comparison tests performed on pooled batches (Panels C, D) or with the nested t-test vs ACTN2wt (Panels F, H, I). Abbreviations: EHTs, engineered heart tissues; N/d, number of EHTs/differentiations.

Similar results were obtained at 1.5 and 2 Hz (data not shown). Overall, ACTN2mut EHTs exhibited a significant force impairment, which can be explained by the marked deficiency of sarcomere-associated proteins detected in 2D-cultured hiPSC-CMs (Table S1) and in EHTs (Table S3).

## 4. Discussion

This study investigated the cellular and functional impacts of an *ACTN2* gene variant (c.740C>T; p.Thr247Met) in hiPSC-CMs. Compared to ACTN2wt, ACTN2mut hiPSC-CMs exhibited (i) cellular hypertrophy, myofibrillar disarray, multinucleation, ACTN2 protein aggregation, and activation of both the UPS and ALP in 2D culture, (ii) a marked reduction in the levels of sarcomere-associated proteins in 2D and EHTs, and iii) force impairment in EHTs. These findings indicate impaired sarcomerogenesis and proteopathy as typical features in ACTN2mut.

We reproduced previous findings observed in 2D-cultured heterozygous ACTN2 (ACTN2het) hiPSC-CMs [8], such as hypertrophy and myofibrillar disarray in ACTN2mut hiPSC-CMs. Furthermore, diseased cells exhibited dysregulation of pathways involved in sarcomere function and proteostasis, and ACTN2mut EHTs exhibited force impairment, resembling a DCM phenotype [27-30]. This is in line with the low abundance of several sarcomeric proteins, including ACTN2 in ACTN2mut hiPSC-CMs, leading to a poorly developed sarcomere and possibly to a more immature cardiomyocyte state. In addition, SYNPO2 and SYNPO2L, which contribute to early assembly and stabilization of the Z-disk via interaction with filamin and ACTN2 [22,27], were also less abundant, supporting disruption of the ACTN2 interactome and deficient sarcomere development in ACTN2mut hiPSC-CMs. The reduced Z-disk integration of exogenous MUT-ACTN2 in ACTN2wt by live cell imaging supports the susceptibility of mutant ACTN2 to aggregate. Conversely, exogenous WT-ACTN2 in ACTN2mut ameliorated sarcomere integration, and partially replaced endogenous mutant without changing the total level of ACTN2. The inverse correlation between sarcomere incorporation and aggregation suggests that non-incorporated mutant proteins form aggregates and contribute to the low level of ACTN2 protein in ACTN2mut hiPSC-CMs. Previous analysis of the dynamic behavior of two *ACTN2* missense variants (p.Ala119Thr and p.Gly111Val), which are also located in the calponin-homology domain, revealed similar phenotypes [31]. Both mutants exhibited reduced binding affinities to F-actin by biochemical assays and alterations of Z-disk localization and dynamic behavior after gene transfer of mEos2-tagged *ACTN2* in adult cardiomyocytes.

The higher levels of several proteins involved in proteostasis such as the UPS and ALP in ACTN2mut hiPSC-CMs found in this study are in agreement with previous findings in HCM septal myectomies [32]. This was associated with a higher chymotrypsin-like activity of the proteasome and global activation of the ALP in ACTN2mut hiPSC-CMs. Even though others have shown that WT ACTN2 is degraded by the UPS [33], the low ACTN2 protein level detected by Western blot and proteomic analysis in the ACTN2mut line was unlikely due to degradation by the UPS or the ALP. This suggests that the global activation of both the UPS and ALP are rather compensatory, protective mechanisms against ACTN2 aggregation. The low abundance of ACTN2 and of several other sarcomeric proteins likely reflects a reduced mRNA translation and protein incorporation into myofilament to maintain the overall stoichiometry of the sarcomere [34]. This might explain the poor and reduced formation of sarcomeres in ACTN2mut hiPSC-CMs. Alteration of Z-disk protein turnover combined with subsequent activation of autophagy has been recently reported in hiPSC-CMs carrying the p.Gly1674X or p.Val1668_Gly1674del *FLNC* variant, resulting in haploinsufficiency and misfolded protein, respectively [35]. Similarly, FLNC protein aggregation and myofibrillar disarray was reported in cardiac muscle specimens of HCM patients carrying the p.Gly2151Ser or p.Ala1539Thr FLNC variant, and was associated with a high risk of sudden cardiac death [36]. These data emphasized the disease-causing role of proteotoxicity in *FLNC*-related cardiomyopathies and presumed its therapeutic potential.

Functional deficits found in ACTN2mut EHTs are in line with a recent disease modeling study that investigated the p.Arg14del variant in phospholamban (PLN; [29]). The authors showed activation of the unfolded protein response as a compensatory, protective mechanism in the setting of PLN-caused hypocontractility in hiPSC-CMs and EHTs. These findings are further supported by the evidence of PLN protein aggregates in a p.Arg14del mouse model [37]. Interestingly, PLN aggregates and altered protein homeostasis pathways were observed before the onset of functional deficits.

To date, only one study has investigated a homozygous truncating *ACTN2* variant (p.Gln860Stop) associated with RCM in mutant carriers [6]. Corresponding hiPSC-CMs displayed hypertrophy, impaired contractility, and myofibrillar disarray. In contrast to our findings, the ACTN2 protein level was not reduced. However, because of the C-terminal truncation, the authors suggested loss of protein-protein interaction as the main cause for disease development. This implies a differing mode of action for truncating and missense *ACTN2* variants, further depending on affected functional domains. Nevertheless, these findings are in line with the diminished contractile function in ACTN2mut EHTs, as the patient affected by the homozygous truncating *ACTN2* variant (p.Gln860Stop) developed RCM and heart failure (HF) at the age of 23 [6]. Based on the severity of cardiomyopathy phenotypes found in this study combined with the recent evidence that *ACTN2* is linked to HF [38], it can be assumed that a patient harboring the homozygous *ACTN2* missense variant would develop DCM or RCM leading to HF.

In conclusion, this study revealed an additional cellular pathology for the p.Thr247Met *ACTN2* variant, leading to proteopathy. Our data indicate the (compensatory) activation of the proteolytic machinery in ACTN2mut hiPSC-CMs, likely to ‘cope’ with protein aggregation.

## 5. Study limitations

When working with hiPSCs it is important to consider possible limitations of the model such as unstable genome integrity, storage of hiPSC lines, immaturity and reproducibility when using hiPSC-CMs (for reviews, see [39-41]). To comply with best cell culture practices, we applied regular karyotyping and genotyping, and established master cell banks of each hiPSC line [42]. However, both ACTN2wt and ACTN2mut hiPSC lines were generated from the ACTN2het hiPSC line with CRISPR/Cas9 genetic tools and homology directed repair (HDR). We found that both hiPSC lines have a correct WT or MUT allele, and that the other recombined allele after HDR contains additional CRISPR/Cas9-mediated on-site defect, i.e. a splicing mutation for ACTN2wt ([19]) and a large genomic rearrangement for ACTN2mut (Figure S1A,B), both leading to nonsense mRNAs (Figure S2D; [19]). Thus, the consequence is the presence of only WT mRNA and protein in the ACTN2wt cardiomyocytes, whereas ACTN2mut cardiomyocytes exhibited only MUT mRNA and protein, making these two cell lines still interesting for comparison. Another limitation is that we cannot decide whether ACTN2 aggregate formation is detrimental, contributing to disease progression, or rather beneficial, to avoid sarcomere incorporation of the mutant protein.

## Supplementary Materials

The following supporting information can be downloaded at: www.mdpi.com/xxx/s1. Figure S1: Generation of ACTN2wt and ACTN2mut cell lines; Figure S2: Validation of cardiac differentiation and mRNA and protein analysis; Figure S3. Representative data of the high-content screening performed in 2D-cultured hiPSC-CMs; Table S1: RNA-seq and mass spectrometry analyses of sarcomere-associated proteins in 2D-cultured ACTN2mut vs. ACTN2wt hiPSC-CMs; Table S2: RNA-seq and mass spectrometry analyses of the UPS and ALP in 2D-cultured ACTN2mut vs. ACTN2wt hiPSC-CMs; Table S3: RNA-seq and mass spectrometry analyses of sarcomere-associated proteins in 3D-cultured ACTN2mut vs. ACTN2wt hiPSC-CMs; Table S4: Acronyms and names of genes evaluated with the nanoString nCounter® Elements technology; Table S5: LC-MS/MS parameter (data dependent mode, spectral library); Table S6: LC-MS/MS parameter (data independent mode; quantitative data); Supplemental experimental procedures: Ethics statement, Generation and culture of hiPSC-derived cardiomyocytes (hiPSC-CMs) in 2D and EHT formats, Southern blot DNA analysis, Immunofluorescence staining of hiPSC-CMs, Morphological analysis of 2D-cultured hiPSC-CMs, Live cell imaging of 2D-cultured hiPSC-CMs, High-content imaging of 2D cultured hiPSC-CMs, RNA isolation and gene expression analysis, Western blot analysis, Measurement of chymotrypsin-like activity of the proteasome, Proteome analysis, Production and purification of adeno-associated virus vector particle; Dataset S1: Mass spectrometry data of 2D ACTN2mut hiPSC-CMs analyzed with IPA; Dataset S2: RNA-seq data of 2D ACTN2mut hiPSC-CMs analyzed with IPA; Video 1: Contracting ACTN2wt hiPSC-CMs transduced with WT-ACTN2 HaloTag; Video 2: Contracting ACTN2wt hiPSC-CMs transduced with MUT-ACTN2 HaloTag; Video 3: Contracting ACTN2wt EHTs. Video 4. Contracting ACTN2mut EHTs.

## Author Contributions

Conceptualization and analysis: ATLZ, MP, LC; Methodology and Investigation: CRISPR/Cas9: MP, NP; Cloning and virus production: ATLZ, SRS, IB; Cardiac differentiation: ATLZ, MP, NP, AM; cell culture, transduction and treatments: ATLZ, MP, MMJ, JB, CSB, AM, GM; EHT generation, maintenance and force measurement: AM; Immunofluorescence and live-cell imaging: ATLZ, MP, JB; Proteomics/bioinformatics: EH, MD, MGS, UV, MP; RNA isolation and gene expression analysis: MP, MDL, EK; RNA-seq: DI, SV; Omics analysis: ATLZ, MP, SC, SV, LC; Western blots: SRS, EO, EA, SaS; High-content imaging: DM; Writing – Original draft: ATLZ, MP and LC; Writing – Review & Editing: ATLZ, MP, SRS, AM, ML, MD, DM, UV, WTP, TE, EH, SaS, LC. All authors reviewed the manuscript and approved the submitted version.

## Institutional Review Board Statement

All procedures were in accordance with the Code of Ethics of the World Medical Association (Declaration of Helsinki). The study was reviewed and approved by the Ethical Committee of the Ärztekammer Hamburg (PV3501).

## Informed Consent Statement

The HCM patient carrying the heterozygous *ACTN2* (c.740C>T; dbSNP ID: rs755492182) mutation was recruited in the outpatient HCM clinic at the University Heart and Vascular Center Hamburg and provided written informed consent for genetic analysis and the use of skin fibroblasts [8].

## Data Availability Statement

Datasets, analysis and study materials will be made available on request to other researchers for purposes of reproducing the results or replicating the procedures. The full description of materials and are provided in the Supplement. All data of OMICs experiments have been made publicly available. The mass spectrometry data have been deposited to the ProteomeXchange Consortium via the PRIDE partner repository with the dataset identifier PXD034258. The RNA-seq data have been deposited to the European Nucleotide Archive (ENA) at EMBL-EBI under accession number PRJEB52889.

## Acknowledgments

The authors gratefully acknowledge Birgit Klampe, Sandra Laufer and Thomas Schulze (Pharmacology, Hamburg) for participating in the production of hiPSC-CMs, Malte Loos for supporting in the hiPSC-CMs treatment and fixation, and the FACS and Microscopy Imaging Facility core facilities (Hamburg). We also thank Boris Skryabin (Transgenic animal and genetic engineering Models (TRAM), Faculty of Medicine of the Westfalian Wilhelms-University, Münster), for the Southern blot analysis and Kerstin Kutsche (Human Genetics, Hamburg) for fruitful discussion. Additionally, we thank Anja Wiechert for support in sample preparation and Stephan Michalik for providing the in-house R pipeline for the quantitative analysis of proteomics data.

