## Supplemental information and dataset for "*ACTN2* mutant causes proteopathy in human iPSC-derived cardiomyocytes": Zech-Prondzynski-Supplemental file.docx

### **Supplement**

#### **Supplemental Figures**

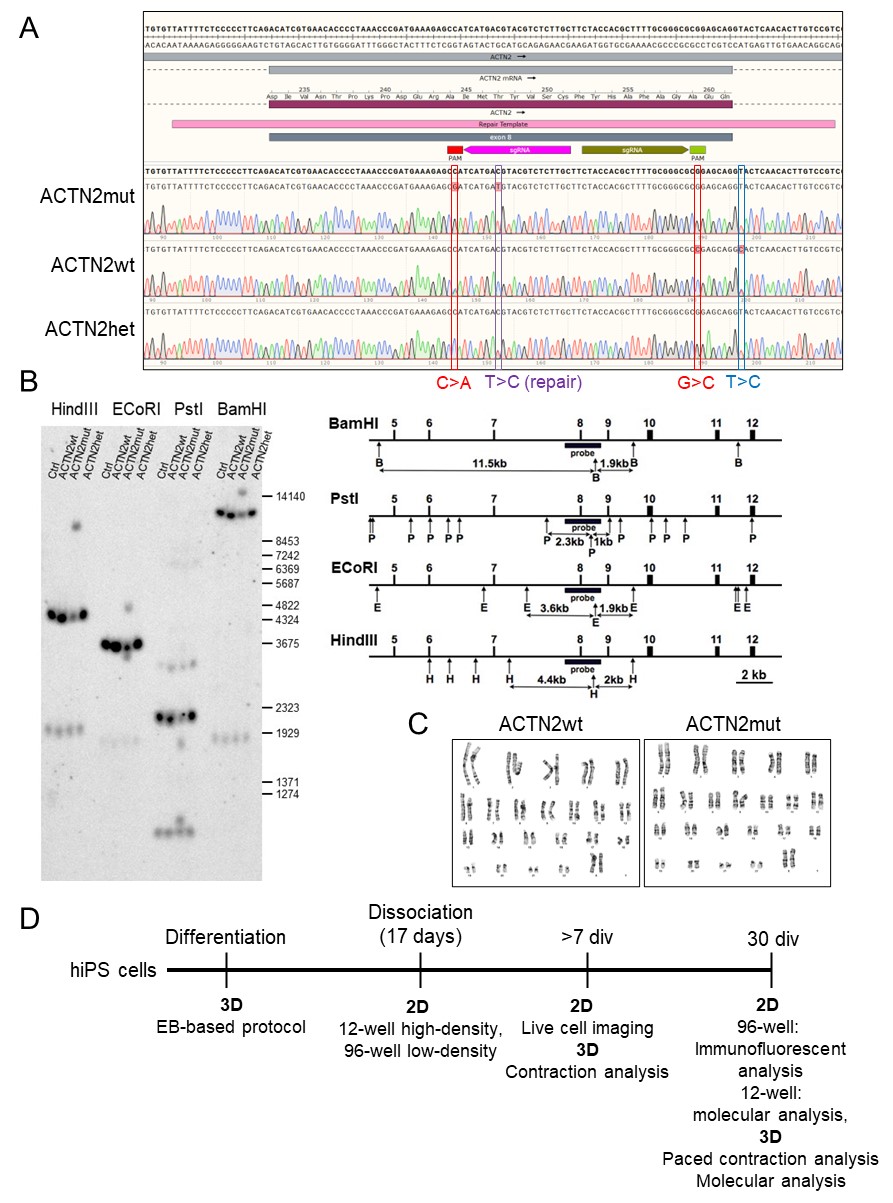

**Figure S1. Generation of the ACTN2wt and ACTN2mut hiPSC lines. (A)** Sanger sequencing results of the affected *ACTN2* locus in exon 8, with the location of the PAM sequences, sgRNAs and repair template used to create the ACTN2wt and ACTN2mut hiPSC lines from the heterozygous ACTN2het hiPSC line. At position 152 is indicated the heterozygous c.740C>T mutation in ACTN2het (purple). ACTN2wt sequence shows the repair at position 152 (p.740C; ACG; pThr247; purple), the two introduced silent mutations (heterozygous G>A at position 144 and homozygous G>C at position 189; red and the on-target effect of CRISPR/Cas at position 197; blue), which induced the skipping of exon 8 and a nonsense mRNA. ACTN2mut sequence shows the mutation at position 152 (p.740T, ATG; pMet247) and no silent mutations, suggesting the presence of only one mutant allele at the locus. **(B)** Southern blot performed with 4 different restriction enzymes on genomic DNA extracted from ACTN2wt, ACTN2mut, ACTN2het and a control (Ctrl) hiPSC line. This reveals a large re-arrangement of one allele in ACTN2mut, affecting the 5’ fragment of all restriction enzymes, leading to a nonsense mRNA. **(C)** G-banding results are depicted by representative karyograms of the investigated hiPSC ACTN2wt (passage 55) and ACTN2wt (passage 68) lines. **(C)** Protocol for the production of hiPSC-derived cardiomyocytes, models and experimental procedures that were used for this study. Abbreviations: div, days in vitro; EB, embryoid body.

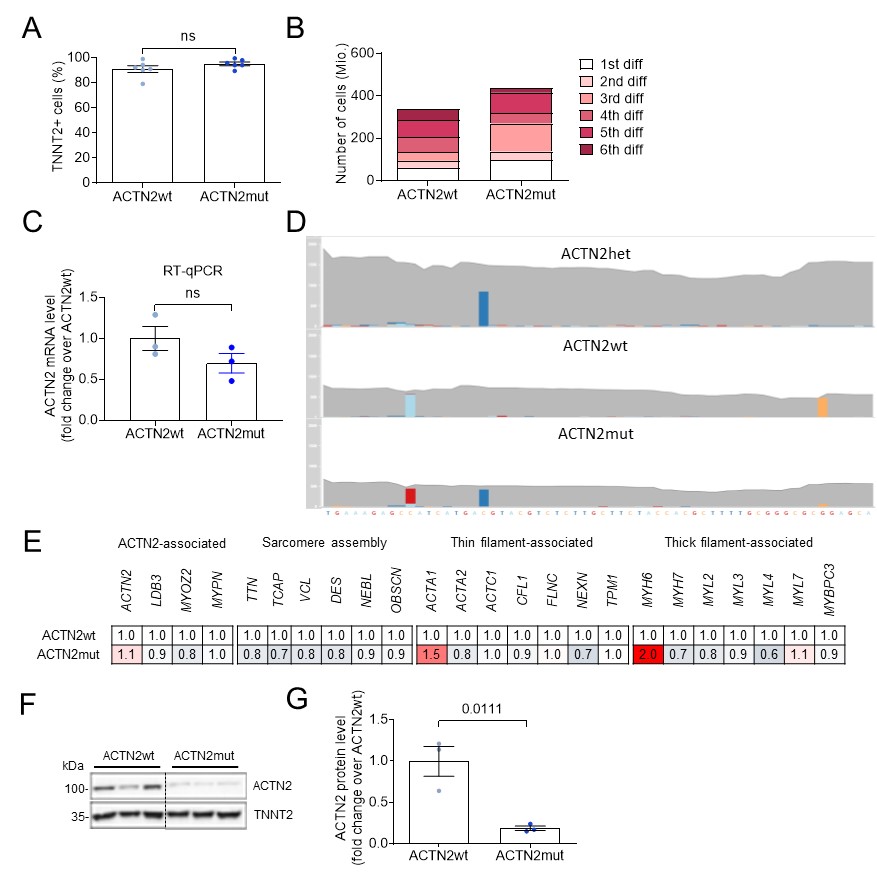

**Figure S2: Validation of cardiac differentiation and mRNA and protein analysis.  (A)** Percentage of TNNT2+ cells in ACTN2wt and ACTN2mut hiPSC-CMs was determined by flow cytometry to validate the quality of cardiac differentiation (n = 6 differentiations). **(B)** Cardiomyocyte yield of six single differentiation runs was evaluated in ACTN2wt and ACTN2mut hiPSC-CMs. **(C)** Quantification of *ACTN2* mRNA by RT-qPCR with the housekeeping gene *GAPDH*. **(D)** RNA-seq of ACTN2 locus showing 2-fold lower RNA counts in both ACTN2wt and ACTN2mut compared to the original heterozygous ACTN2-mutated (ACTN2het; c.740C>T, dark blue) hiPSC line (red to light blue (C>A) and orange (G>C), silent mutations in ACTN2wt). **(E)** Gene expression analysis were performed with the nanoString nCounter® Elements technology. The mRNA counts were normalized to housekeeping genes *ABCF1, CLTC, GAPDH, TUBB* and related to ACTN2wt (ACTN2wt: N/n=18/3, ACTN2mut: N/n=17/3). **(F)** Representative Western is shown performed on SDS-fraction of pooled samples of hiPSC-CMs stained with antibodies against ACTN2 and TNNT2 and **(G)** corresponding quantification of ACTN2 normalized to TNNT2 is shown (ACTN2wt: n/d = 3/1, ACTN2mut: n/d = 3/1). Data are expressed as mean ± SEM, and *P*-values were obtained with the unpaired Student’s t-test. Abbreviations: TNNT2, cardiac troponin T; kDa, kilodalton; N/n/d, number of (pooled) cells/wells/differentiations.

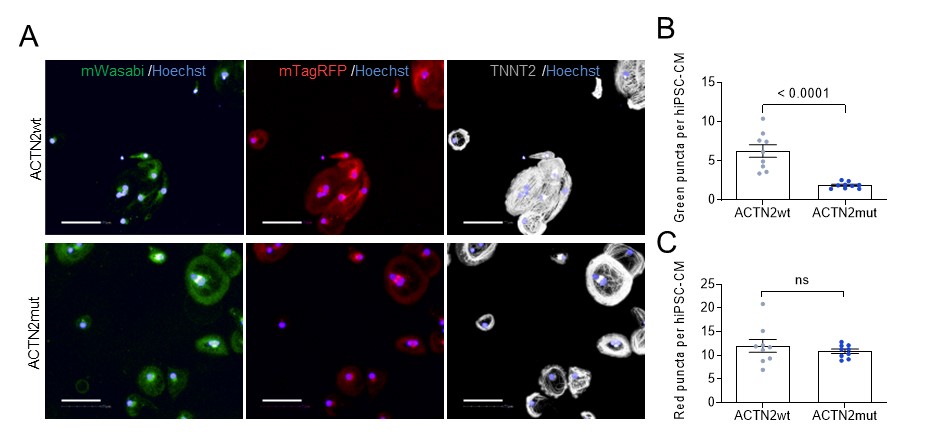

**Figure S3. Representative data of the high-content screening performed in 2D-cultured hiPSC-CMs. (A)** Representative images of the high-content imaging of ACTN2wt and ACTN2mut hiPSC-CMs transduced with an AAV6-mWasabi-mTagRFP-hLC3 tandem construct (MOI 10,000) on day 30 of culture. Subsequently, the hiPSC-CMs were fixed and stained with a TNNT2 (cardiac troponin T) antibody and Hoechst for nuclei staining. Representative images were taken with the Operetta high-content imaging system at 20x magnification (PerkinElmer; Scale bar = 100 µm). **(B)** Number of green puncta and **(C)** number of red puncta per ACTN2wt and ACTN2mut hiPSC-CM (n=wells/treatment per CM per well: ACTN2wt (n=9); ACTN2mut (n=9)). Data are expressed as mean ± SEM, with *P*-values obtained with the unpaired Student’s t-test (panels B, C).

**Supplementary Tables**

**Table S1. RNA-seq and mass spectrometry analyses of sarcomere-associated proteins in 2D-cultured ACTN2mut vs. ACTN2wt hiPSC-CMs.**

| **Cell line** | **Group** | **RNA** | **Log2 ratio** | **FC** | ***P-*value** | **Adjusted *P-*value** | **Group** | **Protein** | **Log2 ratio** | **FC** | ***P-*value** | **Adjusted *P-*value** |
| --- | --- | --- | --- | --- | --- | --- | --- | --- | --- | --- | --- | --- |
| ACTN2mut vs ACTN2wt | Sarcomere-associated | *ACTA1* | 0.84 | 1.78 | 6.98E-02 | 4.42E-01 | Sarcomere-associated | ACTA1 | -0.59 | 0.66 | 6.30E-06 | 2.93E-04 |
|  |  | *ACTC1* | 0.26 | 1.20 | 2.69E-01 | 8.56E-01 |  | ACTC1 | -0.64 | 0.64 | 3.27E-03 | 4.84E-02 |
|  |  | *ACTN1* | -0.26 | 0.84 | 6.57E-01 | 9.83E-01 |  | ACTN1 | 0.48 | 1.39 | 2.63E-05 | 9.58E-04 |
|  |  | *ACTN2* | 0.46 | 1.38 | 2.14E-01 | 8.02E-01 |  | ACTN2 | -1.63 | 0.32 | 3.02E-80 | 2.91E-77 |
|  |  | *DES* | -0.01 | 0.99 | 9.75E-01 | 9.97E-01 |  | DES | 0.31 | 1.24 | 7.03E-02 | 3.34E-01 |
|  |  | *DMD* | -0.40 | 0.76 | 2.54E-01 | 8.44E-01 |  | DMD | -0.21 | 0.87 | 2.10E-02 | 1.70E-01 |
|  |  | *FHL1* | -0.19 | 0.88 | 7.31E-01 | 9.87E-01 |  | FHL1 | 0.56 | 1.47 | 1.34E-02 | 1.30E-01 |
|  |  | *FHL2* | -0.44 | 0.74 | 4.49E-02 | 3.37E-01 |  | FHL2 | 0.27 | 1.20 | 3.74E-02 | 2.37E-01 |
|  |  | *FLNC* | 0.33 | 1.25 | 1.71E-01 | 7.26E-01 |  | FLNC | -0.49 | 0.71 | 3.94E-21 | 1.42E-18 |
|  |  | *JUP* | -0.06 | 0.96 | 8.18E-01 | 9.96E-01 |  | JUP | -0.31 | 0.80 | 5.19E-06 | 2.54E-04 |
|  |  | *LDB3* | 0.21 | 1.15 | 6.18E-01 | 9.80E-01 |  | LDB3 | 0.01 | 1.01 | 1.00E+00 | 1.00E+00 |
|  |  | *MYBPC3* | 0.35 | 1.27 | 3.94E-01 | 9.33E-01 |  | MYBPC3 | -0.82 | 0.57 | 4.61E-51 | 3.32E-48 |
|  |  | *MYH6* | 0.92 | 1.89 | 2.95E-02 | 2.56E-01 |  | MYH6 | -0.52 | 0.70 | 3.35E-02 | 2.23E-01 |
|  |  | *MYH7* | -0.01 | 0.99 | 9.75E-01 | 9.97E-01 |  | MYH7 | -0.89 | 0.54 | 1.23E-10 | 1.54E-08 |
|  |  | *MYL1* | 0.85 | 1.81 | 3.44E-01 | 9.11E-01 |  | MYL1 | -0.81 | 0.57 | 2.17E-115 | 3.13E-112 |
|  |  | *MYL2* | -0.03 | 0.98 | 9.49E-01 | 9.96E-01 |  | MYL2 | -0.65 | 0.64 | 5.23E-03 | 6.92E-02 |
|  |  | *MYL3* | 0.03 | 1.02 | 9.11E-01 | 9.96E-01 |  | MYL3 | -0.68 | 0.62 | 2.10E-09 | 2.33E-07 |
|  |  | *MYL4* | -0.32 | 0.80 | 2.06E-01 | 7.88E-01 |  | MYL4 | -0.58 | 0.67 | 1.31E-06 | 7.15E-05 |
|  |  | *MYOM1* | 0.06 | 1.04 | 8.56E-01 | 9.96E-01 |  | MYOM1 | -0.99 | 0.50 | 1.80E-05 | 7.01E-04 |
|  |  | *MYOZ2* | -0.07 | 0.95 | 8.09E-01 | 9.96E-01 |  | MYOZ2 | -1.00 | 0.50 | 5.66E-42 | 3.26E-39 |
|  |  | *MYPN* | 0.32 | 1.25 | 5.26E-01 | 9.72E-01 |  | MYPN | -0.24 | 0.84 | 1.91E-15 | 4.59E-13 |
|  |  | *NEBL* | -0.14 | 0.90 | 6.14E-01 | 9.80E-01 |  | NEBL | -0.63 | 0.65 | 9.34E-21 | 2.99E-18 |
|  |  | *OBSCN* | 0.15 | 1.11 | 7.25E-01 | 9.87E-01 |  | OBSCN | -0.72 | 0.61 | 8.78E-22 | 3.62E-19 |
|  |  | *OBSL1* | 0.51 | 1.42 | 1.51E-01 | 6.88E-01 |  | OBSL1 | -0.24 | 0.85 | 2.36E-01 | 6.25E-01 |
|  |  | *PKP2* | 0.13 | 1.10 | 6.45E-01 | 9.83E-01 |  | PKP2 | -0.55 | 0.68 | 9.08E-05 | 2.73E-03 |
|  |  | *SLMAP* | -0.34 | 0.79 | 2.04E-01 | 7.84E-01 |  | SLMAP | -0.21 | 0.86 | 9.36E-02 | 3.92E-01 |
|  |  | *SYNPO2* | 0.37 | 1.29 | 4.28E-01 | 9.43E-01 |  | SYNPO2 | -0.40 | 0.76 | 1.78E-02 | 1.54E-01 |
|  |  | *SYNPO2L* | 0.02 | 1.01 | 9.59E-01 | 9.96E-01 |  | SYNPO2L | -0.69 | 0.62 | 3.31E-07 | 2.33E-05 |
|  |  | *TNNC1* | -0.09 | 0.94 | 6.90E-01 | 9.84E-01 |  | TNNC1 | -0.89 | 0.54 | 2.12E-08 | 1.91E-06 |
|  |  | *TNNI1* | 0.44 | 1.36 | 1.17E-01 | 6.03E-01 |  | TNNI1 | -0.87 | 0.55 | 8.09E-07 | 4.67E-05 |
|  |  | *TNNI3* | 0.15 | 1.11 | 5.12E-01 | 9.70E-01 |  | TNNI3 | -0.82 | 0.57 | 4.57E-05 | 1.54E-03 |
|  |  | *TNNT2* | 0.22 | 1.17 | 3.50E-01 | 9.16E-01 |  | TNNT2 | -0.86 | 0.55 | 3.74E-07 | 2.51E-05 |
|  |  | *TPM1* | 0.07 | 1.05 | 7.44E-01 | 9.91E-01 |  | TPM1 | -0.80 | 0.57 | 9.28E-14 | 1.78E-11 |
|  |  | *TPM3* | -0.23 | 0.86 | 6.77E-01 | 9.83E-01 |  | TPM3 | -0.93 | 0.52 | 4.17E-02 | 2.49E-01 |
|  |  | *TTN* | -0.14 | 0.90 | 7.64E-01 | 9.93E-01 |  | TTN | -0.83 | 0.56 | 0.00E+00 | 3.60E-306 |

Purple indicates significantly altered genes or proteins. Abbreviation: FC, Fold change.

###### **Table S2. RNA-seq and mass spectrometry analyses of the UPS and ALP in 2D-cultured ACTN2mut vs. ACTN2wt hiPSC-CMs.**

| **Cell line** | **Group** | **RNA** | **Log2 ratio** | **FC** | ***P-*value** | **Adjusted *P-*value** | **Group** | **Protein** | **Log2 ratio** | **FC** | ***P-*value** | **Adjusted *P-*value** |
| --- | --- | --- | --- | --- | --- | --- | --- | --- | --- | --- | --- | --- |
| ACTN2mut vs ACTN2wt | UPS and ALP | *BNIP3* | 1.85 | 3.61 | 0.0006 | 6.16E-03 | UPS and ALP | ASAH1 | 0.89 | 1.85 | 2.29E-05 | 8.68E-04 |
|  |  | *CTSH* | 1.13 | 2.18 | 0.0000 | 6.25E-05 |  | BAG3 | 0.38 | 1.30 | 2.30E-02 | 1.79E-01 |
|  |  | *DEPTOR* | -1.74 | 0.30 | 0.0016 | 1.75E-02 |  | CTSC | 0.67 | 1.59 | 1.39E-03 | 2.61E-02 |
|  |  | *KLHL13* | -1.35 | 0.39 | 0.0000 | 1.93E-04 |  | CTSL | 1.07 | 2.10 | 2.26E-03 | 3.76E-02 |
|  |  | *MAP1LC3A* | -0.94 | 0.52 | 0.0246 | 2.23E-01 |  | FAM98A | 1.38 | 2.61 | 1.68E-02 | 1.50E-01 |
|  |  | *MDM2* | -0.95 | 0.52 | 0.0002 | 1.32E-03 |  | GBA | 0.38 | 1.30 | 4.71E-02 | 2.69E-01 |
|  |  | *STAT1* | -1.06 | 0.48 | 0.0463 | 3.44E-01 |  | HSPA1A | 0.23 | 1.18 | 2.37E-03 | 3.89E-02 |
|  |  | *TRIM4* | -6.35 | 0.01 | 0.0000 | 4.38E-07 |  | PSMA3 | 0.38 | 1.30 | 1.97E-02 | 1.62E-01 |
|  |  |  |  |  |  |  |  | PSMA6 | 0.37 | 1.29 | 6.47E-04 | 1.47E-02 |
|  |  |  |  |  |  |  |  | PSMB5 | 0.45 | 1.37 | 3.68E-02 | 2.35E-01 |
|  |  |  |  |  |  |  |  | PSMC6 | 0.29 | 1.22 | 2.02E-03 | 3.42E-02 |
|  |  |  |  |  |  |  |  | PSMD11 | 0.31 | 1.24 | 1.11E-03 | 2.15E-02 |
|  |  |  |  |  |  |  |  | PSME1 | 0.46 | 1.37 | 5.36E-06 | 2.54E-04 |
|  |  |  |  |  |  |  |  | PSME2 | 0.38 | 1.30 | 3.78E-02 | 2.39E-01 |
|  |  |  |  |  |  |  |  | TRIM54 | 0.44 | 1.36 | 2.60E-03 | 4.10E-02 |
|  |  |  |  |  |  |  |  | UBA1 | 0.39 | 1.31 | 7.83E-07 | 4.67E-05 |
|  |  |  |  |  |  |  |  | UBE2O | 0.68 | 1.60 | 2.53E-02 | 1.88E-01 |
|  |  |  |  |  |  |  |  | UBQLN2 | 0.70 | 1.62 | 7.99E-04 | 1.70E-02 |

Purple indicates significantly altered genes or proteins. Abbreviation: FC, Fold change.

**Table S3. RNA-seq and mass spectrometry analyses of sarcomere-associated proteins in 3D-cultured ACTN2mut vs. ACTN2wt hiPSC-CMs (EHTs).**

| **Cell line** | **Group** | **RNA** | **Log2 ratio** | **FC** | ***P-*value** | **Adjusted *P-*value** | **Group** | **Protein** | **Log2 ratio** | **FC** | ***P-*value** | **Adjusted *P-*value** |
| --- | --- | --- | --- | --- | --- | --- | --- | --- | --- | --- | --- | --- |
| ACTN2mut vs ACTN2wt | Sarcomere-associated | *ACTA1* | 2.79 | 6.91 | 6.13E-05 | 9.47E-02 | Sarcomere-associated | ACTA1 | -0.55 | 0.69 | 1.30E-04 | 1.63E-02 |
|  |  | *ACTC1* | 0.71 | 1.64 | 2.41E-04 | 1.74E-01 |  | ACTC1 | -0.47 | 0.72 | 6.47E-02 | 1.00E+00 |
|  |  | *ACTN1* | 0.38 | 1.30 | 1.18E-01 | 9.04E-01 |  | ACTN1 | 0.03 | 1.02 | 1.00E+00 | 1.00E+00 |
|  |  | *ACTN2* | 0.10 | 1.07 | 6.47E-01 | 9.94E-01 |  | ACTN2 | -1.41 | 0.38 | 1.14E-56 | 1.65E-53 |
|  |  | *DES* | 0.61 | 1.53 | 1.41E-02 | 6.46E-01 |  | DES | -0.42 | 0.75 | 8.26E-07 | 1.40E-04 |
|  |  | *DMD* | -0.14 | 0.91 | 5.65E-01 | 9.91E-01 |  | DMD | -0.15 | 0.90 | 1.00E+00 | 1.00E+00 |
|  |  | *FHL1* | -0.04 | 0.97 | 9.49E-01 | 9.99E-01 |  | FHL1 | -0.43 | 0.74 | 2.81E-01 | 1.00E+00 |
|  |  | *FHL2* | 0.09 | 1.06 | 7.03E-01 | 9.95E-01 |  | FHL2 | -0.14 | 0.91 | 8.59E-01 | 1.00E+00 |
|  |  | *FLNC* | 0.70 | 1.63 | 2.72E-02 | 7.50E-01 |  | FLNC | -0.09 | 0.94 | 1.00E+00 | 1.00E+00 |
|  |  | *JUP* | -0.31 | 0.81 | 9.16E-02 | 8.80E-01 |  | JUP | -0.63 | 0.65 | 2.91E-12 | 1.20E-09 |
|  |  | *LDB3* | -0.07 | 0.95 | 6.90E-01 | 9.95E-01 |  | LDB3 | -0.44 | 0.74 | 7.13E-05 | 1.00E-02 |
|  |  | *MYBPC3* | 0.20 | 1.15 | 2.87E-01 | 9.70E-01 |  | MYBPC3 | -0.94 | 0.52 | 9.00E-49 | 8.65E-46 |
|  |  | *MYH6* | 0.43 | 1.35 | 1.52E-01 | 9.23E-01 |  | MYH6 | -0.29 | 0.82 | 3.75E-03 | 2.17E-01 |
|  |  | *MYH7* | 0.04 | 1.03 | 8.86E-01 | 9.97E-01 |  | MYH7 | -0.95 | 0.52 | 1.16E-01 | 1.00E+00 |
|  |  | *MYL1* | 1.11 | 2.15 | 3.21E-01 | 9.75E-01 |  | MYL1 | -0.74 | 0.60 | 7.06E-94 | 2.04E-90 |
|  |  | *MYL2* | -0.18 | 0.88 | 5.62E-01 | 9.90E-01 |  | MYL2 | -0.84 | 0.56 | 1.55E-02 | 5.40E-01 |
|  |  | *MYL3* | -0.35 | 0.78 | 1.18E-01 | 9.04E-01 |  | MYL3 | -0.69 | 0.62 | 1.54E-08 | 4.92E-06 |
|  |  | *MYL4* | -0.29 | 0.82 | 1.69E-01 | 9.36E-01 |  | MYL4 | -0.56 | 0.68 | 1.14E-07 | 2.52E-05 |
|  |  | *MYOM1* | 0.28 | 1.21 | 1.79E-01 | 9.38E-01 |  | MYOM1 | -1.01 | 0.50 | 5.87E-04 | 5.46E-02 |
|  |  | *MYOZ2* | -0.56 | 0.68 | 1.47E-03 | 3.39E-01 |  | MYOZ2 | -1.01 | 0.50 | 3.52E-39 | 2.54E-36 |
|  |  | *MYPN* | -0.01 | 0.99 | 9.55E-01 | 9.99E-01 |  | MYPN | -0.56 | 0.68 | 3.72E-11 | 1.34E-08 |
|  |  | *NEBL* | -0.12 | 0.92 | 5.48E-01 | 9.90E-01 |  | NEBL | -0.54 | 0.69 | 8.10E-07 | 1.40E-04 |
|  |  | *OBSCN* | 0.22 | 1.17 | 2.42E-01 | 9.64E-01 |  | OBSCN | -0.35 | 0.79 | 3.44E-04 | 3.54E-02 |
|  |  | *OBSL1* | 0.29 | 1.23 | 9.31E-02 | 8.81E-01 |  | OBSL1 | -0.06 | 0.96 | 1.00E+00 | 1.00E+00 |
|  |  | *PKP2* | -0.37 | 0.78 | 6.96E-02 | 8.44E-01 |  | PKP2 | -0.65 | 0.64 | 1.70E-08 | 4.92E-06 |
|  |  | *SLMAP* | -0.10 | 0.94 | 6.47E-01 | 9.94E-01 |  | SLMAP | -0.39 | 0.76 | 1.08E-01 | 1.00E+00 |
|  |  | *SYNPO2* | -0.51 | 0.70 | 1.78E-01 | 9.38E-01 |  | SYNPO2 | -0.79 | 0.58 | 4.08E-06 | 6.19E-04 |
|  |  | *SYNPO2L* | 0.04 | 1.03 | 8.38E-01 | 9.97E-01 |  | SYNPO2L | -0.76 | 0.59 | 1.54E-07 | 3.17E-05 |
|  |  | *TNNC1* | -0.53 | 0.69 | 1.42E-02 | 6.46E-01 |  | TNNC1 | -0.87 | 0.55 | 1.13E-06 | 1.81E-04 |
|  |  | *TNNI1* | -0.02 | 0.99 | 9.22E-01 | 9.99E-01 |  | TNNI1 | -0.81 | 0.57 | 3.52E-07 | 6.77E-05 |
|  |  | *TNNI3* | -0.19 | 0.88 | 5.84E-01 | 9.92E-01 |  | TNNI3 | -0.75 | 0.60 | 4.00E-03 | 2.26E-01 |
|  |  | *TNNT2* | -0.13 | 0.91 | 4.76E-01 | 9.88E-01 |  | TNNT2 | -0.84 | 0.56 | 1.04E-07 | 2.51E-05 |
|  |  | *TPM1* | -0.17 | 0.89 | 3.68E-01 | 9.75E-01 |  | TPM1 | -0.70 | 0.62 | 8.46E-13 | 4.07E-10 |
|  |  | *TPM3* | -0.41 | 0.75 | 7.99E-02 | 8.62E-01 |  | TPM3 | -0.80 | 0.58 | 6.25E-01 | 1.00E+00 |
|  |  | *TTN* | 0.01 | 1.01 | 9.53E-01 | 9.99E-01 |  | TTN | -0.05 | 0.96 | 1.00E+00 | 1.00E+00 |

Purple indicates significantly altered genes or proteins. Abbreviation: FC, Fold change.

###### **Table S4. Acronyms and names of genes evaluated with the nanoString nCounter® Elements technology.**

| **Gene** | **Acronym** | **Accession number (NCBI)** |
| --- | --- | --- |
| Alpha-Actinin 2 | *ACTN2* | NM_001103.2 |
| LIM Domain Binding 3 | *LDB3* | NM_001080116.1 |
| Myozenin 2 | *MYOZ2* | NM_016599.4 |
| Myopalladin | *MYPN* | NM_032578.2 |
| Titin | *TTN* | NM_133432.1 |
| Titin-cap | *TCAP* | NM_003673.3 |
| Vinculin | *VCL* | NM_014000.2 |
| Desmin | *DES* | NM_001927.3 |
| Nebulin | *NEBL* | NM_001173484.1 |
| Obscurin | *OBSCN* | NM_001098623.2 |
| Actin alpha 1, skeletal muscle | *ACTA1* | NM_001100.3 |
| Actin Alpha 2, smooth muscle | *ACTA2* | NM_001613.1 |
| Actin alpha, cardiac muscle 1 | *ACTC1* | NM_005159.4 |
| Cofilin 1 | *CFL1* | NM_005507.2 |
| Filamin C | *FLNC* | NM_001127487.1 |
| Nexilin | *NEXN* | NM_144573.3 |
| Tropomyosin 1 | *TPM1* | NM_000366.5 |
| Myosin heavy chain 6 | *MYH6* | NM_002471.3 |
| Myosin heavy chain 7 | *MYH7* | NM_000257.2 |
| Myosin Regulatory Light Chain 2 | *MYL2* | NM_000432.3 |
| Myosin Regulatory Light Chain 3 | *MYL3* | NM_000258.2 |
| Myosin Regulatory Light Chain 4 | *MYL4* | NM_002476.2 |
| Myosin Regulatory Light Chain 7 | *MYL7* | NM_021223.2 |
| Cardiac myosin binding protein C | *MYBPC3* | NM_000256.3 |
| ATP Binding Cassette Subfamily F Member 1 | *ABCF1* | NM_001090.2 |
| Clathrin Heavy Chain | *CLTC* | NM_004859.2 |
| Glyceraldehyde 3-phosphate dehydrogenase | *GAPDH* | NM_002046.3 |
| Tubulin Beta Class I | *TUBB* | NM_178014.3 |

###### **Table S5. LC-MS/MS parameter (data dependent mode, spectral library).**

| ***Data dependent analyses (DDA)*** |  |
| --- | --- |
| ***Reversed phase liquid chromatography*** | **Ultimate 3000 RSLC (Thermo Scientific)** |
| *Trap column* | 75 μm inner diameter, packed with 3 μm C18 particles (Acclaim PepMap100, Thermo Scientific) |
| *Analytical column* | Accucore^TM^ 150-C18, 25 cm x 75 μm, 2.6 μm C18, 150 Å (Thermo Scientific) |
| *Flow rate* | 300 nl/min |
| *Column oven temperature* | 40 °C |
| *Buffer system* | Binary buffer system consisting of 0.1% acetic acid in HPLC-grade water (buffer A) and 100% ACN in 0.1% acetic acid (buffer B) |
| *Gradient* | Gradient of buffer B: 2 min 2% to 5 %, 10 min 5%, 120 min 5% to 25%, 5 min 25% to 40%, 2 min 40% to 90%, 5 min 90%, 3 min 90% to 2%, 10 min 2% |
| ***Mass spectrometer*** | **Q Exactive HFx** |
| *Operation mode* | Data-dependent |
| *Electrospray* | Nanospray Flex Ion Source |
| ***Full MS*** |  |
| *MS scan resolution* | 60,000 |
| *AGC target* | 3.00E+06 |
| *Maximum ion injection time for the MS scan* | 45 ms |
| *Scan range* | 350 to 1650 *m/z* |
| *Spectra data type* | Profile |
| ***dd-MS2*** |  |
| *Resolution* | 15,000 |
| *MS/MS AGC target* | 1.00E+05 |
| *Maximum ion injection time for the MS/MS scans* | 22 ms |
| *Spectra data type* | Profile |
| *Selection for MS/MS* | 12 most abundant isotope patterns with charge ≥2 from the survey scan |
| *Isolation window* | 1,3 *m/z* |
| *Fixed first mass* | 100 *m/z* |
| *Dissociation mode* | HCD |
| *Normalized collision energy* | 27% |
| *Dynamic exclusion* | 45 s |
| *Charge exclusion* | 1,>6 |

###### **Table S6. LC-MS/MS parameter (data independent mode; quantitative data).**

| ***Data independent analyses (DIA)*** |  |
| --- | --- |
| ***Reversed phase liquid chromatography*** | **Ultimate 3000 RSLC (Thermo Scientific)** |
| *Trap column* | 75 μm inner diameter, packed with 3 μm C18 particles (Acclaim PepMap100, Thermo Scientific) |
| *Analytical column* | Accucore^TM^ 150-C18, 25 cm x 75 μm, 2.6 μm C18, 150 Å (Thermo Scientific) |
| *Flow rate* | 300 nl/min |
| *Column oven temperature* | 40 °C |
| *Buffer system* | Binary buffer system consisting of 0.1% acetic acid in HPLC-grade water (buffer A) and 100% ACN in 0.1% acetic acid (buffer B) |
| *Gradient* | Dradient of buffer B: 2 min 22% to 5 %, 8 min 5%, 120 min 5% to 25%, 5 min 25 to 40%, 2 min 40% to 90%, 5 min 90%, 3 min 90% to 2%, 10 min 2% |
| ***Mass spectrometer*** | **Q Exactive HFx** |
| *Operation mode* | Data-independent |
| *Electrospray* | Nanospray Flex Ion Source |
| ***Full MS*** |  |
| *MS scan resolution* | 120,000 |
| *AGC target* | 3.00E+06 |
| *Maximum ion injection time for the MS scan* | 60 ms |
| *Scan range* | 350 to 1200 *m/z* |
| *Spectra data type* | Profile |
| ***DIA MS2*** |  |
| *Resolution* | 30,000 |
| *MS/MS AGC target* | 3.00E+06 |
| *Maximum ion injection time for the MS/MS scans* | Auto |
| *Spectra data type* | Profile |
| *selection for MS/MS* | 1 |
| *Isolation window* | 70 windows *m/z* 10, 11 *m/z* |
| *Fixed first mass* | 200 |
| *Dissociation mode* | Higher energy collisional dissociation (HCD) |
| *Normalized collision energy* | 27.5% |
| *Dissociation mode* | HCD |

### **Video legends**

**Video 1. Contracting ACTN2wt hiPSC-CMs transduced with WT-*ACTN2* HaloTag.** Widefield fluorescence microscopy of contracting ACTN2wt hiPSC-CMs that were transduced with AAV6-*TNNT2*-WT-*ACTN2*-HaloTag and stained for imaging with TMR-ligand to visualize sarcomeres.

**Video 2. Contracting ACTN2wt hiPSC-CMs transduced with MUT-*ACTN2* HaloTag.** Widefield fluorescence microscopy of contracting ACTN2wt hiPSC-CMs that were transduced with AAV6-*TNNT2*-MUT-*ACTN2*-HaloTag and stained for imaging with TMR-ligand to visualize sarcomeres.

**Video 3. Contracting ACTN2wt EHTs.** Widefield video microscopy of contracting ACTN2wt EHTs that were paced at 1 Hz to analyze functional parameters independent of varying baseline frequencies.

**Video 4. Contracting ACTN2mut EHTs.** Widefield video microscopy of contracting ACTN2mut EHTs that were paced at 1 Hz to analyze functional parameters independent of varying baseline frequencies.

### **Dataset legends**

**Dataset S1. MS data of 2D ACTN2mut hiPSC-CMs analyzed with IPA.** Significantly altered proteins were submitted to an unsupervised analysis of ACTN2mut in relation to ACTN2wt hiPSC-CMs. Fisher’s exact test, unadjusted *P*-value <0.05.

**Dataset S2. RNA-seq data of 2D ACTN2mut hiPSC-CMs analyzed with IPA.** Significantly altered protein-coding genes were submitted to an unsupervised analysis of ACTN2mut in relation to ACTN2wt hiPSC-CMs. Fisher’s exact test, unadjusted *P*-value <0.05.

#### **Supplemental experimental procedures**

**Ethics statement**

The HCM patient carrying the *ACTN2* (c.740C>T; dbSNP ID: rs755492182) mutation was recruited in the outpatient HCM clinic at the University Heart and Vascular Center Hamburg and provided written informed consent for genetic analysis and the use of skin fibroblasts [1]. All procedures were in accordance with the Code of Ethics of the World Medical Association (Declaration of Helsinki). The study was reviewed and approved by the Ethical Committee of the Ärztekammer Hamburg (PV3501).

**Generation and culture of hiPSC-derived cardiomyocytes (hiPSC-CMs) in 2D and EHT formats**

Generation of both wild-type (ACTN2wt) and mutant (ACTN2mut) hiPSC lines was performed with CRISPR/Cas9 gene tools in ACTN2het hiPSC [1]. Both clones were analyzed for the same off-targets as published previously. Additionally, PCR fragments of cells containing the modified *ACTN2* locus were subcloned using the CloneJET PCR Cloning Kit (Thermo Scientific, Vilnius, Lithuania) for discrimination of allele-specific genotypes. Finally, all used cell lines were genotyped by PCR for the *ACTN2* locus and karyotyped by G-banding as reported previously [1]. To amplify specific DNA fragments, PCR was conducted according to standard protocols using PrimeSTAR® HS DNA Polymerase (Takara, Kusatsu, Japan) and AmpliTaq® Gold DNA polymerase (Thermo Fisher Scientific, Vilnius, Lithuania) following manufacturer’s instructions.

Cardiomyocyte (CM) differentiation from hiPSCs was performed following a three-step protocol with generation of embryoid bodies (EBs) in spinner flasks and cardiac differentiation efficiency was determined by flow cytometry as described previously [2]. After dissociation with collagenase 2 (200 units/mL, LS004176, Worthington, Lakewood, NJ) hiPSC-CMs were plated on Geltrex-coated (1:100, Gibco, Grand Island, NY) 96-well or 12-well plates at a density of 2,500 - 5,000 cells/well or 440,000 cells/well, respectively. HiPSC-CMs were maintained in culture for 7 days in 96-well plates (µclear®, Greiner Bio-One, Frickenhausen, Germany) when used for live cell imaging and for 30 days for immunofluorescent analysis, as well as 12-well plates, which were used for molecular analysis, at 37 °C in 7% CO_2_ and atmospheric O_2_ (21%) prior to further analysis. Furthermore, single cell suspensions of hiPSC-CMs were subjected to the 3D format of EHTs, which were generated in a 24-well format (800,000 hiPSC-CM/EHT) in a fibrin matrix, maintained in culture and analyzed from day 26 on as described previously [1].

**Southern blot DNA analysis**

Southern blot was performed as previously described [3]. HiPSC pellets were lysed in buffer containing 100 mM tris-HCl (pH 8.5), 5 mM EDTA, 0.2% SDS, 200 mM NaCl, and proteinase K (100 μg/ml; Roche) overnight at 55 °C. Genomic DNA was extracted by phenol-chloroform and chloroform, followed by precipitation with 2.5 volumes of isopropanol and washing with 70% ethanol. The DNA pellet was dissolved in TE buffer [10 mM tris (pH 7.9) and 0.2 mM EDTA]. Approximately 10 to 20 μg of genomic DNA was digested with the corresponding restriction endonuclease, fractionated on 0.8% agarose gels, and transferred to GeneScreen nylon membranes (NEN DuPont). The membranes were hybridized with ^32^P-labeled specific DNA probe. DNA labeling was performed using a random prime DNA labeling kit (Roche) and [α-^32^P] deoxycytidine-5′ triphosphate (PerkinElmer). Membranes were washed with 0.5x saline sodium phosphate EDTA (SSPE) buffer [1× saline sodium phosphate EDTA buffer is 0.18 M NaCl, 10 mM NaH2PO4, and 1 mM EDTA (pH 7.7)] and 0.5% SDS at 65 °C and exposed to MS film (Kodak) at −80 °C.

##### **Immunofluorescence staining of hiPSC-CMs**

HiPSC-CMs were cultured for 30 days in 96-well plates (µclear®, Greiner Bio-One, Frickenhausen, Germany) and subsequently prepared for immunofluorescence analysis as described previously [1]. The following primary antibodies were used: ACTN2 – A7811 (1:800), Sigma-Aldrich, St.Louis, MO; cardiac troponin T (TNNT2) – ab8295 (1:200) or ab45932 (1:500), Abcam, Cambridge, UK; HaloTag – G9281 (1:400), Promega, Madison, WI. The following secondary antibodies were used: anti-mouse Alexa Fluor® 488 antibody - LT A11029 (1:800), Life Technologies, Eugene, OR; anti-rabbit Alexa Fluor® 546 antibody - LT A11035 (1:800), Life Technologies, Eugene, OR; anti-rabbit Alexa Fluor® 647 – LT A11031 (1:800), Life Technologies, Eugene, OR. Nuclei staining was obtained with Hoechst 33342 (1 µg/mL; Thermo Fisher Scientific, Waltham, MA). The Wheat Germ Agglutinin (WGA) Alexa Fluor® 633 conjugate (W21404, 1:50, Thermo Fisher Scientific, Waltham, MA) was used to perform the volume measurement of hiPSC-CMs. Images were obtained by confocal microscopy using a Zeiss LSM 800 confocal microscope with a 40x-oil immersion objective.

##### **Morphological analysis of 2D-cultured hiPSC-CMs**

Analysis of ACTN2 protein aggregates was performed using ImageJ. Therefore, immunofluorescence images were imported to ImageJ and subjected to the “find Maxima” tool adjusting the signal threshold to 254 and 1 in fixed hiPSC-CMs and live-cell imaging experiments, respectively. The automatically generated image was then analyzed with the “find particles” tool, and areas of ACTN2 aggregates were automatically determined by ImageJ. Finally, ACTN2 areas were normalized to a control line or condition to express differences in areas as fold changes.

The volume measurement of hiPSC-CMs was performed using Imaris 7.6.1. software. Z-stack images were imported to Imaris and a surface was added. After choosing default parameters, the relevant channel (WGA) was selected. Next, a smooth surface was selected (surface area detail level of 0.312 µm), and the threshold was set with the absolute intensity. Subsequently, the volume (µm³) of the generated surface was calculated.

##### **Live cell imaging of 2D-cultured hiPSC-CMs**

After cardiac differentiation, hiPSC-CMs were transduced for 1 h in suspension at 37 °C with an adeno-associated virus serotype 6 (AAV6) carrying either the wild-type (WT; *ACTN2*-WT-HaloTag®) or the mutant (MUT; *ACTN2*-MUT-HaloTag®) construct under the control of the human *TNNT2* promoter at a MOI of 10,000 [4]. Ultimately, transduced hiPSC-CMs were seeded in Geltrex-coated (1:100; Gibco, Grand Island, NY) 96-well plates at a density of 2,500 - 5,000 cells per well. Live-cell imaging experiments were performed after 7 days in culture by adding TMR-ligand to the medium (0.1 mM; G8251, Promega, Madison, WI) for 30 min, specifically staining ACTN2-HaloTag® protein, followed by washing twice with culture medium and 10 min of Hoechst 33342 incubation (1 µg/mL; H3570, Thermo Fisher Scientific, Waltham, MA), which was diluted in culture medium, followed by a final washing step. Images of hiPSC-CMs were acquired by confocal microscopy using a Zeiss LSM 800 confocal microscope, whereby only beating hiPSC-CMs were included to ensure cell viability. Quantification of integration events was carried out in a blinded fashion and expressed as percentage of transduced hiPSC-CMs.

##### **High-content imaging of 2D cultured hiPSC-CMs**

HiPSC-CMs were either gently thawed or dissociated at day 17 into single cells using collagenase II [2] and transduced with an AAV6-*TNNT2*-mTagRFP-mWasabi-hLC3 (human LC3) tandem construct (MOI 10,000) for 1 h at 37 °C, while inverting the tube every 10 min, including a non-transduced control. HiPSC-CMs were seeded at a density of 2,500 cells in 96-well plates. After 30 days of culture, hiPSC-CMs were washed twice with PBS and fixed with Roti-Histofix (Roth, Karlsruhe, Germany) for 20 min at 4 °C. After three additional washing steps with PBS, hiPSC-CMs were stained with a TNNT2 antibody (1:500; ab45932) and with Hoechst 33342 (1 µg/mL; Thermo Fisher Scientific, Waltham, MA), as previously described.[1,5] Automated image acquisition was performed in the Operetta high-content imaging system (Perkin Elmer), by capturing mWasabi signal (green), mTagRFP signal (red) and counterstained nuclei (Hoechst) and TNNT2 (far-red), at 20x magnification. Since mWasabi is quenched in the acidic autolysosomes, green puncta were considered as autophagosomes. Assuming the red puncta to represent autophagosomes plus lysosomes, we then quantified the red puncta minus green puncta as autolysosomes in hiPSC-CMs. This was done by Harmony high-content imaging analysis software, by repurposing previously developed algorithms for determination of cardiomyocyte-relevant markers [6]. In brief, hiPSC-CMs were identified by displaying positive TNNT2 signal and quantification of green and red puncta (number and intensity) was done in the whole cell area using Harmony in-built functions.

##### **RNA isolation and gene expression analysis**

Total RNA was extracted from 2D-cultured hiPSC-CMs using TRIzol® Reagent (Life Technologies) following the manufacturer's protocol. For expression analysis with the nanoString nCounter® Elements technology a total amount of 50 ng RNA was hybridized with a customized nanoString Gene Expression CodeSet (Table S4 in the Data Supplement) and analyzed using the nCounter® Sprint Profiler (nanoString). The mRNA levels were determined with the nSolver™ Data Analysis Software including background subtraction using negative controls and normalization to housekeeping genes (*ABCF1, CLTC, GAPDH, TUBB;* Table S4 in the Data Supplement) and expressed as fold change over ACTN2wt for the respective groups. For RNA sequencing analysis, samples derived from 2-3 independent differentiations were analyzed for each comparison. Therefore, total RNA was extracted from individual 2D cell pellet or EHT (2-3 biological replicates per differentiation) and after concentration measurement, equal amounts of RNA from 2-3 hiPSC-CMs in 2D-cultured or EHT format of each biological replicate (differentiation) were pooled and prepared as previously reported [1].

For RNA-seq analysis, gene abundance were quantified using salmon (v0.12.0) [7], and human gene annotations from gencode (version 33) for GRCh38 genome assembly were imported using R package tximport [8]. Counts normalization and differential gene expression analysis was performed using DESeq2 package [9]. Null variance of Wald test statistic output by DESeq2 was re-estimated using R package fdrtool [10] to calculate *P*-values (and adjusted using Benjamini-Hochburg method) for the final list of differentially expressed genes. FDR (BH-adjusted *P*-value) < 0.1 and log2 fold change < X was used as criteria for the final DE gene list.

##### **Western blot analysis**

Protein extraction from 2D-cultured hiPSC-CMs was based on a recent publication [11] with adjusted volumes as follow. For protein extraction from untreated 2D-cultured hiPSC-CMs, 50 µL water with protease inhibitor cocktail was added to the cell pellet (about 440,000 cells/well) to obtain the cytosolic fraction. Subsequently, the membrane-enriched fraction was collected by adding 75 µL SDS-buffer to the pellet. For protein extraction from treated 2D-cultured hiPSC-CMs, 100 µL SDS-buffer was added to the cell pellet, the samples homogenized and spun down for 15 min at 14 000 rpm at RT. The supernatant was kept has total crude protein lysate. Western blot analysis was performed according to previous publications [12,13] with slight adaptations. For analysis of 30-day-old, 2D-cultured hiPSC-CMs, total crude protein lysates were evaluated by mixing 7.5 µg of protein with 6x Laemmli buffer and subsequent separations on 4–15% Mini-PROTEAN Precast Gels (Bio-Rad, Hercules, CA) or 10%/12% self-casted acrylamide/bisacrylamide (29:1) gels. After electrophoresis, proteins were transferred to nitrocellulose or PVDF membranes and stained with the following antibodies: ACTN2 – A7811(1:10,000), Sigma-Aldrich, St. Louis, MO; TNNT2 – ab8295 (1:5000), Abcam, Cambridge, UK; GAPDH – 5G4 (1:5000), HyTest, Turku, Finland; LC3B – 2775 (1:1000), Cell Signaling Technology, Danvers, MA; SQSTM1 – P0067 (1:2000), Sigma-Aldrich, St. Louis, MO; ubiquitin – FK2 clone, BML-PW8810 (1:10,000), Enzo Life Sciences, Farmingdale, NY. Signals were detected using the SuperSignal West Dura Extended Duration Substrate (Thermo Fisher Scientific, Rockford, IL), acquired with the ChemiDoc Touch Imaging System (Bio-Rad, Hercules, CA) and quantified with the Image Lab software (Bio-Rad, Hercules, CA).

##### **Measurement of chymotrypsin-like activity of the proteasome**

##### Cell pellet (about 440,000 cells/well) of 2D-cultured hiPSC-CMs was dissolved in 50 µL water with protease inhibitor cocktail (complete miniTM, Roche Diagnostics, Rotkreuz, Switzerland). After three freeze-thaw-cycles the cell lysate was homogenized by pipetting up and down and centrifuged at 4 °C, full speed for 30 min in a table-top centrifuge. The supernatant was kept as the cytosolic fraction. The measurement of the chymotrypsin-like activity of the proteasome was based on previous publications [11,14] and was slightly adapted for the evaluation in the cytosolic fraction of untreated 30-day-old, 2D-cultured hiPSC-CMs. To determine the activity of the 20S proteasome, 10 µg of protein were used for the measurement which was performed in the absence of ATP and with a final concentration of 60 µM of the synthetic fluorogenic substrate Suc-LLVY-AMC (Enzo Life Sciences, BML-P802).

##### **Proteome analysis**

*Generation of protein samples:* Samples derived from 3 independent differentiations were analyzed for each comparison. Initially, protein was extracted from individual 2D cell pellet or EHT (2-3 biological replicates per differentiation) by five cycles of freezing (liquid nitrogen) and thawing (30 °C, 1,400 rpm) in 8 M urea/2 M thiourea. Cell debris and insoluble material was separated by centrifugation (20,000 × g, 1 h at 4 °C). After determination of protein content with a Bradford assay (Biorad, Munich, Germany), equal protein amounts from 2-3 hiPSC-CMs in 2D culture or EHT format of each biological replicate (differentiation) were pooled and subjected to proteolytic digestion.

*Sample preparation:* 4 µg of total protein from each sample were reduced (2.5 mM DTT ultrapure, Invitrogen, for 15 min at 37 °C) and alkylated (10 mM iodacetamide, Sigma Aldrich, for 30 min at 37 °C). Proteolysis was performed using LysC (1:100 for 3 h at 37 °C), followed by tryptic digestion overnight at 37 °C (both from Promega, Madison, WI, USA). The tryptic digestion was stopped by adding acetic acid (final concentration 1%) followed by desalting using ZipTip-µC18 tips (Merck Millipore, Darmstadt, Germany). Eluted peptides were concentrated by evaporation under vacuum and subsequently resolved in 0.1% acetic acid, 2% acetonitrile (ACN) containing HRM/iRT peptides (Biognosys, Zurich, Switzerland) according to manufacturer’s recommendation.

*Mass Spectrometry Measurements:* Mass spectrometric (MS) data was recorded on a QExactive HFx mass spectrometer (Thermo Electron, Bremen, Germany). Before MS data acquisition tryptic peptides were separated by reverse phase chromatography (Accucore^TM^ 150-C18, 25 cm x 75 μm, 2.6 μm C18, 150 Å) using an Ultimate 3000 nano-LC system (Thermo Scientific, Waltham, MA, USA) at a constant temperature of 40 °C and a flow rate of 300 nL/min.

To design a spectral library, MS/MS peptides were separated by 2 or 3 h-linear gradients with increasing acetonitrile concentration from 5 to 25% in 0.1% acetic acid, and data were recorded in data dependent mode (DDA). The MS scans were carried out in a *m/z* range of 350 to 1650 *m*/*z*. Data were acquired with a resolution of 60,000 and an AGC target of 3 x 10^6^ at maximal injection times of 45 ms. The top 12 most abundant isotope patterns with charge ≥2 from the survey scan were selected for fragmentation by high energy collisional dissociation (HCD) with a maximal injection time of 22 ms, an isolation window of 1.3 *m/z*, and a normalized collision energy of 27.5 eV. Dynamic exclusion was set to 45 s. The MS/MS scans had a resolution of 15,000 and an AGC target of 1 x 10^5^.

The acquisition of MS data for relative quantitation was performed in data independent mode (DIA) after peptide pre-fractionation using a 120 min-linear gradient from 5% to 25% acetonitrile in 0.1% acetic acid. Briefly, the data was acquired in the *m*/*z* range from 350 to 1200 *m*/*z*, the resolution for MS was 120,000 and for MS/MS 30,000. The AGC target was 3 x 10^6^ for MS and MS/MS. The number of DIA isolation windows was 70 of 11 *m/z* with 1 *m*/*z* overlap. For further details to the instrumental setup and the parameters for LC-MS/MS analysis in DDA and DIA mode see Table S5 and Table S6 in the Data Supplement.

*Data analysis:* Proteins were identified using Spectronaut^TM^ Pulsar 13.4 software (Biognosys AG) against a spectral library generated from data-dependent acquisition measurements of individual and pooled samples of the study. The spectral library construction by Spectronaut was based on a database search using a human protein database (Uniprot vs 03_2019, 20404 entries) extended by sequences of bovine fibrinogen subunits A, B, and C. Because of the use of horse serum as medium supplement, sequences of 10 proteins reproducibly identified by proteotypic peptides were added to the database. The target-decoy search was performed with a parent mass error of ±20 ppm, fragment mass error of 0.01 Da, and allowing full-tryptic peptides (trypsin/P cleavage rule) with a minimal peptide length of 6 amino acids and up to two internal cleavage sites. The search included carbamidomethylation at cysteine as fixed modification and oxidation at methionine and acetylation at protein N-termini as variable modifications. The generation of the ion library in Spectronaut^TM^ v13.4.190802.43655 resulted in a constructed library consisting of 335,310 fragments, 30,756 peptides and 3,376 protein groups.

The Spectronaut DIA-MS analysis was carried out as described previously[15] with project specific modifications. Briefly, the following parameter settings were applied: dynamic MS1 and MS2 mass tolerance, dynamic XIC RT extraction window, automatic calibration, dynamic decoy strategy (library size factor = 0.1, minimum limit = 5000), protein Q-value cutoff of 0.01, precursor Q-value cutoff of 0.001. The search included variable and static modifications as described above for spectral library construction. A local cross run normalization was performed using complete profiles with a Q-value < 0.001. The MS2 peak area was quantified and reported. Missing values were parsed using an iRT profiling strategy with carry-over of exact peak boundaries (minimum Q-value row selection = 0.001). Only non-identified precursors were parsed with a Q-value > 0.0001. Ion values were parsed when at least 20% of the samples contained high quality measured values. Peptides were assigned to protein groups and protein inference was resolved by the automatic workflow implemented in Spectronaut. Only proteins with at least two identified peptides were considered for further analyses.

Data has been median normalized on ion level before statistical analysis was carried out on peptide level after exclusion of peptides with oxidized methionine using the algorithm ROPECA.[16] Binary differences have been identified by application of a moderate t-test.[17] Multiple test correction was performed according to Benjamini and Hochberg.[18] Variance within the data set was visualized by principal component analyses and differences in the protein pattern by Volcano plots.

##### **Production and purification of adeno-associated virus vector particle**

To generate the AAV transfer plasmid pFBGR-TNNT2-GFP, a PCR was performed (primer pair: 5’-GAG CGG CCG CAC GCG TCT CAG TCC ATT AGG AGC CAG TAG C-3’ and 5’-GGG CGA ATT GGG TAC CTT ACT TGT ACA GCT CGT CCA TGC CG-3’) and inserted into linearized pFBGR-GFP using the In-Fusion HD Cloning Kit (639648, Takara Bio Europe SAS, Kusatsu, Japan) according to the manufacturer’s instructions.

The basis of the here applied mTagRFP-mWasabi-hLC3 tandem construct was kindly provided by Dr Zhou [19], and we modified and subcloned it in our lab.^56^ Therefore, the vector pBudCE4.1-CMV (Invitrogen, Carlsbad, CA) was digested with Bam*H*I and HindIII. Touchdown PCR was performed using AmpliTaqGold (Thermo Fisher Scientific, Vilnius, Lithuania) and plasmid pcDNA3.1-mTagRFP-mWasabi-hLC3 as template with the following primer pair: 5’-TTC GAC AAG CTT ACC ATG AGC GAG CTG ATT AAG GAG AAC ATG CAC ATG A-3’ and 5’-TTC GAC GGA TCC TTA CAC TGA CAA TTT CAT CCC GAA CGT CTC CTG GGA GG-3’. Ligation of the PCR product and the digested vector pBudCE4.1-CMV was performed using T4 Ligase according to manufacturer’s instructions (Thermo Fisher Scientific, Vilnius, Lithuania). Subsequently, heat transformation using competent *E. coli* bacteria was performed. In short, bacteria were thawed on ice, 50 ng of DNA (ligation approach) was added, the tube once flicked and incubated for 30 min on ice. The heat shock was performed at 42 °C for 45 sec, followed by a 5-min incubation on ice. After the addition of 200 µL S.O.C. medium, the approach was incubated for 1 h at 37 °C gently shaking, before being plated on ampicillin agar plates. DNA of selected clones was isolated according to manufacturer’s instructions (NucleoSpin Plasmid Mini kit, Macherey-Nagel, Düren, Germany) and evaluated by restriction digest and sequencing.

To express the mTagRFP-mWasabi-hLC3 tandem construct under control of the *TNNT2* promoter with an ampicillin resistance and the ß-Globin IgG intron for higher transcription efficiency, the pGG2-*TNNT2*-insert plasmid was digested with Bam*H*I and NheI. Touchdown PCR was performed using AmpliTaqGold and plasmid pBudCE4.1-CMV-mTagRFP-mWasabi-hLC3 as template with the primer pair: 5’-TTC GAC GGA TCC TTA CAC TGA CAA TTT CAT CCC GAA CGT CTC CTG GGA GG-3’ and 5’-TTC GAC GCT AGC GCC ACC ATG AGC GAG CTG ATT AAG G-3’. Ligation of the PCR product and the digested pGG2-*TNNT2*-insert plasmid, heat transformation using competent *E. coli* bacteria and evaluation of selected clones by restriction digest and sequencing was performed as described above. For virus production, the pGG2-*TNNT2-*mTagRFP-mWasabi-hLC3 was inserted into pFBGR-*TNNT2* by In-Fusion cloning according to manufacturer's instructions. mTagRFP-mWasabi-hLC3 was amplified from pGG2-*TNNT2*-mTagRFP-mWasabi-hLC3 using PrimeStar GXL Polymerase (Takara Bio Europe SAS, Kusatsu, Japan) using 5’-AAC ATC GAT TGA ATT CAC CAT GAG CGA GCT GAT TAA GG-3’ and 5’-TAT AGG GCG AAT TGG GTA CCT TAC ACT GAC AAT TTC ATC CCG A-3’. pFBGR-Ultra-*TNNT2*-GFP was cut with EcoR*I* and Kpn*I* and both fragments were inserted simultaneously generating the AAV transfer plasmid pFBGR-Ultra-*TNNT2*-mTagRFP-mWasabi-hLC3.

For live-cell imaging experiments, WT- or MUT-*ACTN2* and HaloTag were assembled and inserted into pFBGR under control of a *TNNT2* promoter by In-Fusion cloning. Two fragments were generated by PCR using PrimeStar GXL Polymerase (Takara Bio Europe SAS, Kusatsu, Japan): *ACTN2* was amplified from plasmids either containing the WT or Mut sequence (c.740C>T; p.Thr247Met) that were kindly provided by Mathias Gautel using 5’-AAC ATC GAT TGA ATT CAC CAT GAA CCA GAT AGA GCC CGG-3’ and 5’-GCT CGA GGA CAG ATCG CTC TCC CCG TAG AG-3’. HaloTag was amplified from the vector pFC14K-HaloTag®-CMV-Flexi® (G9651, Promega, Madison, WI) using 5’-GAT CTG TCC TCG AGC GAG GAT CTG TAC TTT CAG AGC GAT AAC G-3’ and 5’-GGG CGA ATT GGG TAC CTT AAC CGG AAA TCT CCA GAG TAG ACA GC-3’. pFBGR-Ultra-*TNNT2*-GFP was digested with Eco*R*I and KpnI and both fragments were inserted simultaneously generating the AAV transfer plasmid pFBGR-Ultra-*TNNT2*-*ACTN2*-WT-/MUT-HaloTag. Final AAV transfer plasmids were confirmed by restriction digestion and sequencing. Recombinant baculovirus DNA carrying AAV genomic components was verified by PCR using TNNT2/T7 promoter specific primers.

AAV were produced according to the “titerless infected-cells preservation and scale-up (TIPS)” also known as baculovirus-infected insect cells (BIIC).[20] Sf9 cells were transfected with recombinant baculovirus DNA derived from pFBGR-Ultra-TNNT2-mTagRFP-mWasabi-hLC3 or pFBGR-Ultra-*TNNT2-ACTN2*-WT-/MUT-HaloTag and pSR646-Ultra mCherry using TransIT®-Insect Transfection Reagent (MoBITec, Goettingen, Germany) according to the manufacturer’s instructions. After 3 days, Sf9 cells (Novagen, Darmstadt, Germany) were collected, and 2E+07 fresh Sf9 cells were infected with this primary stock. On the next day, vital fluorescing TriEx Sf9 Cells (Novagen, Darmstadt, Germany) were collected and frozen at 2E+06 cells BIIC stocks in InsectXpress medium (Biozym Scientific GmbH, Verviers, Belgium)/10% DMSO. For AAV production, ~2E+08 fresh Sf9 cells were cultivated in InsectXpress Medium at a density of 2E+06 cells/ml. ~2E+06 Sf9 BIIC stocks containing the expression cassette TNNT2-mTagRFP-mWasabi-hLC3 or TNNT2-WT-/MUT-*ACTN2*-HaloTag and ~2E+06 Sf9 BIIC stocks containing the desired AAV packaging elements were added. AVV lysis buffer (50 mM Tris base, 100 M NaCl, 5 mM MgCl_2_, pH 8.5) was used for harvest. To remove cell debris, centrifugation for 20 min at 12,000 x g was performed, and vector containing lysates were purified using Iodixanol (OptiPrep, PROGEN Biotechnik, Heidelberg, Germany) step gradients. 40% Iodixanol layers were harvested, and iodixanol was removed by ultrafiltration (Amicon Ultra-4, PLQK Ultracel-PL Membran, 50 kDa, MerckMillipore, Darmstadt, Germany).
